# Global mapping of *Salmonella enterica*-host protein-protein interactions during infection

**DOI:** 10.1101/2020.05.04.075937

**Authors:** Philipp Walch, Joel Selkrig, Leigh A. Knodler, Mandy Rettel, Frank Stein, Keith Fernandez, Cristina Viéitez, Clément M. Potel, Karoline Scholzen, Matthias Geyer, Klemens Rottner, Olivia Steele-Mortimer, Mikhail M. Savitski, David W. Holden, Athanasios Typas

## Abstract

Intracellular bacterial pathogens inject effector proteins into host cells to hijack diverse cellular processes and promote their survival and proliferation. To systematically map effector-host protein-protein interactions (PPIs) during infection, we generated a library of 32 *Salmonella enterica* serovar Typhimurium (*S*Tm) strains expressing chromosomally encoded affinity-tagged effector proteins, and quantified PPIs in macrophages and epithelial cells by Affinity-Purification Quantitative Mass-Spectrometry. Thereby, we identified 25 previously described and 421 novel effector-host PPIs. While effectors converged on the same host cellular processes, most had multiple targets, which often differed between cell types. Using reciprocal co-immunoprecipitations, we validated 13 out of 22 new PPIs. We then used this host-pathogen physical interactome resource to demonstrate that SseJ and SseL collaborate in redirecting cholesterol to the *Salmonella* Containing Vacuole (SCV) via NPC1, PipB directly recruits the organelle contact site protein PDZD8 to the SCV, and SteC promotes actin bundling by directly phosphorylating formin-like proteins.

## Introduction

To usurp host defenses, pathogens produce and secrete proteins that directly intercept and modify the endogenous host cell machinery. For intracellular pathogens, this becomes even more important, as they need to actively evade detection by cytoplasmic host innate immune receptors and establish a favorable intracellular niche to ensure their proliferation (Cunha and Zamboni 2013). In turn, the host has evolved mechanisms to overcome such molecular insults. This evolutionary arms race has driven many pathogens to develop remarkably diverse arsenals of effector proteins, as in the case of the bacterial pathogen *Legionella pneumophila* which secretes >300 effectors (Schroeder 2017). Host-pathogen protein-protein interactions (PPIs) are thereby manifold and play a pivotal role in shaping infection outcomes.

Discovering the host targets of effectors has traditionally been the first step to investigate the role of single effectors in infection. The development of methodologies for global PPI profiling in single organisms has also opened the doors for systematically mapping host-pathogen interfaces (Shah et al. 2015). Both global yeast two-hybrid studies (Uetz et al. 2006; Blasche et al. 2014; Calderwood et al. 2007; Shapira et al. 2009) and affinity-tag purification/mass spectrometry (AP/MS) screens (Jäger, Cimermancic, et al. 2011; Penn et al. 2018; Sontag et al. 2016; D’Costa et al. 2019) have been employed to systematically map PPIs at the bacterial- and viral-host interfaces. Initial global PPI efforts often resulted in high false-positive rates in the identification of effector interaction partners and generated skepticism in the community for such studies (Stynen et al. 2012; Rajagopala, Hughes, and Uetz 2009). However, as methodologies and data analysis advanced, large-scale studies are now playing a more active role in resolving the picture of relevant PPIs at the host-pathogen interface. One such case constitutes HIV infection, where more than a thousand PPIs had been reported in literature for just a handful of viral proteins, based on targeted approaches (Jäger, Gulbahce, et al. 2011). Systematic AP-MS resolved the picture, identifying the strong and relevant physical interactions (Jäger, Cimermancic, et al. 2011) and fueled a plethora of mechanistic insights into HIV biology (Chou et al. 2013; Jäger, Kim, et al. 2011). Despite their power, such studies are still limited in their capacity to faithfully recapitulate the infection environment. Until now, PPIs have typically been probed within mammalian cells in which a single effector is overexpressed at a time, in the absence of the pathogen, or by using *in vitro* setups where lysates are passed through columns with immobilized effectors. Besides using non-physiological levels of the effector, such experiments also poorly reflect the infection state *in vivo* due to the absence of infection-relevant rewiring of the host proteome and the presence of additional effectors, which may promote or hinder interactions. Therefore, methods that probe host-pathogen PPIs in the infection context are still in high demand.

To identify effector-host PPIs in their native infection context, we developed a proteomics-based methodology to extract *Salmonella enterica* serovar Typhimurium (*S*Tm)-delivered effectors directly from infected cells and quantify their interacting protein partners. Although *S*Tm is perhaps the best-studied intracellular bacterial pathogen, we still lack a good understanding of the 34 known effectors that are translocated by its two T3SS, with less than half of them having known host targets (Ramos-Morales 2012; Schleker et al. 2012; LaRock, Chaudhary, and Miller 2015; Jennings, Thurston, and Holden 2017). We constructed a library of 32 chromosomally-tagged effectors translocated into the host cytoplasm by both T3SS1 (encoded on *Salmonella* pathogenicity island 1 (SPI-1)) and T3SS2 (encoded on SPI-2) (Jennings, Thurston, and Holden 2017; Ramos-Morales 2012), and used it to profile effector-

host PPIs across two different, relevant cell lines, HeLa and RAW264.7. Thereby, we were able to reconstruct the most comprehensive *S*Tm-host interactome to date, spanning a total of 15 effectors and 421 novel PPIs, and displaying a high degree of intracellular connectivity. The accuracy of this resource was verified by the detection of 25 previously described PPIs, as well as by validating novel interactions using reciprocal pulldowns. Network analysis revealed that diverse effectors targeted host proteins with related functions, with several effectors converging on the same process, and in some cases even interacting. Despite this, most effectors had multiple targets, often in unrelated host cellular processes. Whereas several PPIs were detected in both cell lines tested, most PPIs were specific to the cellular context. Capitalizing on this resource, we further resolve the effector interplay between SseJ and SseL in cholesterol trafficking, demonstrate that PipB directly recruits the endoplasmic reticulum (ER) tethered protein PDZD8 to the *S*Tm-containing vacuole (SCV), and discover that the effector kinase SteC promotes actin bundling via interactions with formin-like proteins (FMNL). Overall, we provide a new method for probing host-pathogen PPIs in a physiological context, and a rich resource that can be used for the discovery of novel *S*Tm infection mechanisms.

## Results

### Affinity-purification quantitative mass-spectrometry (AP-QMS) for mapping the host targets of *Salmonella* effectors during infection

To systematically map the PPI landscape between *S*Tm effectors and mammalian host targets, we generated a library of 32 tagged-effector *S*Tm 14028s strains, i.e. nearly all the known effector proteins translocated by T3SS1 and T3SS2 (see Table S1). To mimic the infection context and ensure physiological effector dosage to the host-cell cytoplasm *via* an active T3SS, we introduced an in-frame C-terminal Strep(2x)-TEV-FLAG(3x) (STF) tag onto the endogenous chromosomal locus of the effector. One exception to this cloning strategy was SifA, where a C-terminal STF tag would otherwise have inactivated the prenylation motif (Reinicke et al. 2005). In this case, we inserted the STF tag into an internal site known to preserve SifA function (Brumell, Goosney, and Finlay 2002) using a two-step cloning process (see Experimental Procedures). Strains expressing chromosomally tagged effectors were then tested for effector expression and translocation to the host cytoplasm during infection of epithelial or macrophage cells, two relevant cell types for *S*Tm infection (LaRock, Chaudhary, and Miller 2015). As expected, effector expression and translocation were most robustly detected at later stages of infection (Figure S1), when intracellular *S*Tm loads were high. We detected a total of 20 effectors (2 from T3SS1, 12 from T3SS2 and 6 from both) being injected at significant levels into the Tx-100 soluble fraction of host cells – this fraction contains the host cytoplasm and organelles, but not the nucleus and intact *S*Tm. These 20 effectors were then used in large-scale infections for AP-QMS analysis (Figure 1A, Table S1).

**Figure 1.**
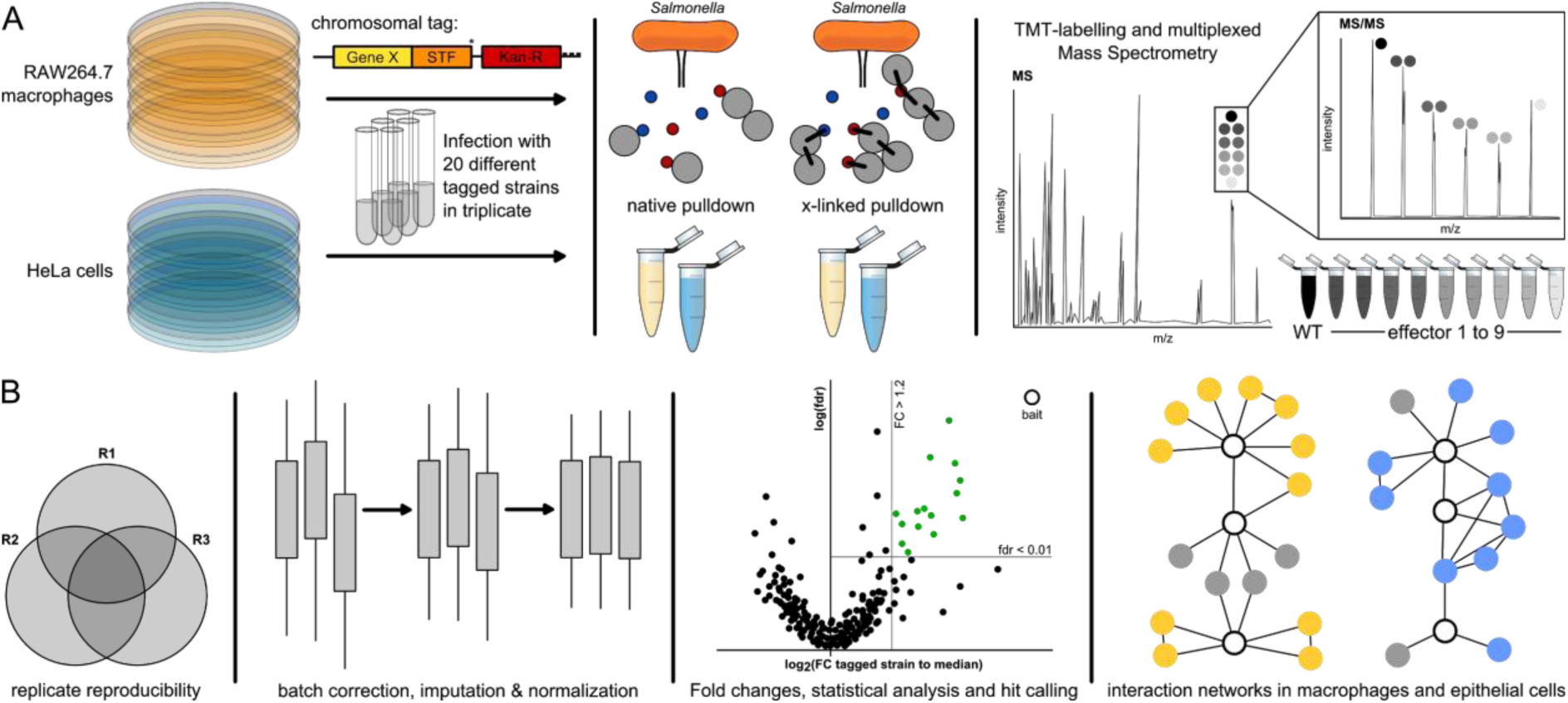
AP-QMS pipeline for mapping effector-host protein interactions during *Salmonella* infection. (A) *Salmonella enterica* Typhimurium 14028s (*S*Tm) strains engineered to express C-terminally tagged effectors with STF – FLAG(2x)-TEV-STREP(3x), or internally STF-tagged SifA, were used to infect HeLa and RAW264.7 cells at a MOI ∼100:1 in three biological replicate experiments. At 20 hpi, cells were washed with PBS, half of the samples were treated with DSP crosslinker for 2h and lysed, and the other half were directly lysed. Lysates containing injected effectors were used for anti-FLAG pulldown and competitively eluted with FLAG peptide. Crosslinked samples were quenched prior to harvesting. Eluates from the pulldowns were reduced, alkylated, cleaned up, digested by trypsin and combined in a TMT-10plex labelling run. We combined elutions from nine different STF-tagged effectors and one untagged wildtype background control (see Experimental Procedures for more information). (B) Only proteins quantified with at least two unique peptides and identified in at least two out of the three biological replicates were used in analysis. Data was checked for reproducibility between replicates (Figure S2A); batch effects were removed using the Limma package, variance was normalized and missing values were imputed (see Experimental Procedures). Differential expression was calculated with respect to the median of the replicate (Figure S2B). A protein was annotated as a ‘hit’ when the false discovery rate (fdr) was < 1% and exhibited a fold increase of at least 20%. We further refined this list by loosening the fdr requirement to < 5% if a PPI passed the FC requirement in both conditions (native and crosslinked). Subsequently, only the strongest 20 PPIs per effector with respect to FC or fdr, as well as PPIs detected in both the native and crosslinked pulldowns, were kept for the final hit list. All analyzed data, or hits only, are listed in Table S2 and S3, respectively. Volcano plots of all pulldowns can be found in Figures S3-6. PPI networks were built from hits passing the above thresholds and known host functional interactions.

To be able to compare our dataset with previous global *S*Tm-host PPI studies (Sontag et al. 2016; D’Costa et al. 2019) and targeted studies (summarized in (LaRock, Chaudhary, and Miller 2015; Jennings, Thurston, and Holden 2017)), we tested PPIs in two commonly used cell lines for *S*Tm infections: HeLa and RAW264.7, which are of distinct cellular and organismal origin (human epithelial and murine macrophages, respectively). We performed FLAG-immunoprecipitation at 20 hours post infection (hpi) under both native (for stable interactions) and cross-linking conditions (for transient interactions) using the cell permeable and reducible cross-linker DSP (Figure 1A). To ensure reproducible quantification of bait and prey proteins relative to background, pulldown eluates were combined in groups of 10 (layout consisting of 9 distinct effector pulldown eluates and one untagged background control) and analyzed in a single 10-plex Tandem Mass Tag (TMT (Werner et al. 2014)) in biological triplicates (Figure 1A). Only proteins identified with at least two unique peptides and found in at least two biological replicates were used for further analysis (Figure 1B). We verified replicate reproducibility (Figure S2A), corrected batch effects, imputed missing values between runs and normalized the median values across each run to ensure accurate sample comparison (Experimental Procedures; Figure 1B). We calculated specific protein enrichment by comparing protein abundance (signal sum) in each TMT channel relative to the median abundance (signal sum) within each TMT10 run for each protein (Figure 1B), which was more robust than comparing to the untagged background strain (Figure S2B), and displayed data as volcano plots (Figures S3-S6). The entire dataset for both cell lines is summarized in Table S2. We detected the bait protein for 13 effectors in both RAW264.7 and HeLa cells, with significant interactions for 12 effectors in RAW264.7 and 9 in HeLa cells. Due to the 20 hpi time point, T3SS2 effectors were, as expected, more readily detected. The resulting hits for each bait (fold change (FC) ≥ 1.2; False Discovery Rate (fdr) ≤ 0.01, after adjusting stringency for hits in both native and cross-linked conditions and capping the number of hits per effector; see Experimental Procedures) are reported in Table S3 and were used to build PPI networks (Figure 1B).

Across the 2 cell lines and 15 effectors, we detected 462 non-redundant PPIs. Of these, 446 PPIs were effector-target, 15 were the baits themselves and 1 was a clear contaminant (IgG-heavy chain). Of the 446 effector-target PPIs; 421 were effector-host (25 previously reported (Table S4), and 396 new) and 25 were effector-bacterial protein interactions. Of those effectors where PPIs were detected, on average, each effector had 19.7 PPIs in RAW macrophages and 26.4 PPIs in HeLa cells. This suggests that the majority of effectors display promiscuous protein-binding inside host cells. Overall, our AP-QMS method robustly captures previously observed *S*Tm effector-host PPIs, while identifying many new ones.

### *Salmonella* effectors target diverse host processes in macrophages and epithelial cells

Using the significant interactions we detected by AP-QMS, and known human, murine or bacterial protein functional interactions (Table S5, STRING DB version 11 (Szklarczyk et al. 2019)), we built two separate PPI networks in RAW264.7 and HeLa cells (Figures 2A and 3A; Experimental Procedures). The networks contained a number of previously characterized PPIs, such as SseJ directly interacting with the host Rho GTPase proteins RhoA and RhoB (Ohlson et al. 2008) in RAW264.7 and HeLa cells (RhoB was not detected even as background in HeLa cells, likely due to low abundance), but the majority of interactions reported were new (Figures 2B and 3B). In total, we detected 25 previously reported interactions (Table S4): e.g. PipB2-KLC1/2, PipB2-KIF5B, SseL-OSBP and SseI-ACADM (Sontag et al. 2016; Henry et al. 2006; Auweter et al. 2012) in the two cell lines. We failed to capture some well-described PPIs, such as that of SifA-SKIP (Jackson et al. 2008; Diacovich et al. 2009; Zhao et al. 2015) or AvrA-MKK7 (Jones et al. 2008; Du and Galán 2009). False negatives are common in AP-MS protocols (Verschueren et al. 2015) and can have multiple causes (see Discussion). In addition, several of the new interactions may be indirect and mediated *via* another host protein (piggybacking is a common issue of AP approaches; (Nesvizhskii 2012; Teng et al. 2015)), which would explain effectors binding to multiple host proteins of the same process.

**Figure 2.**
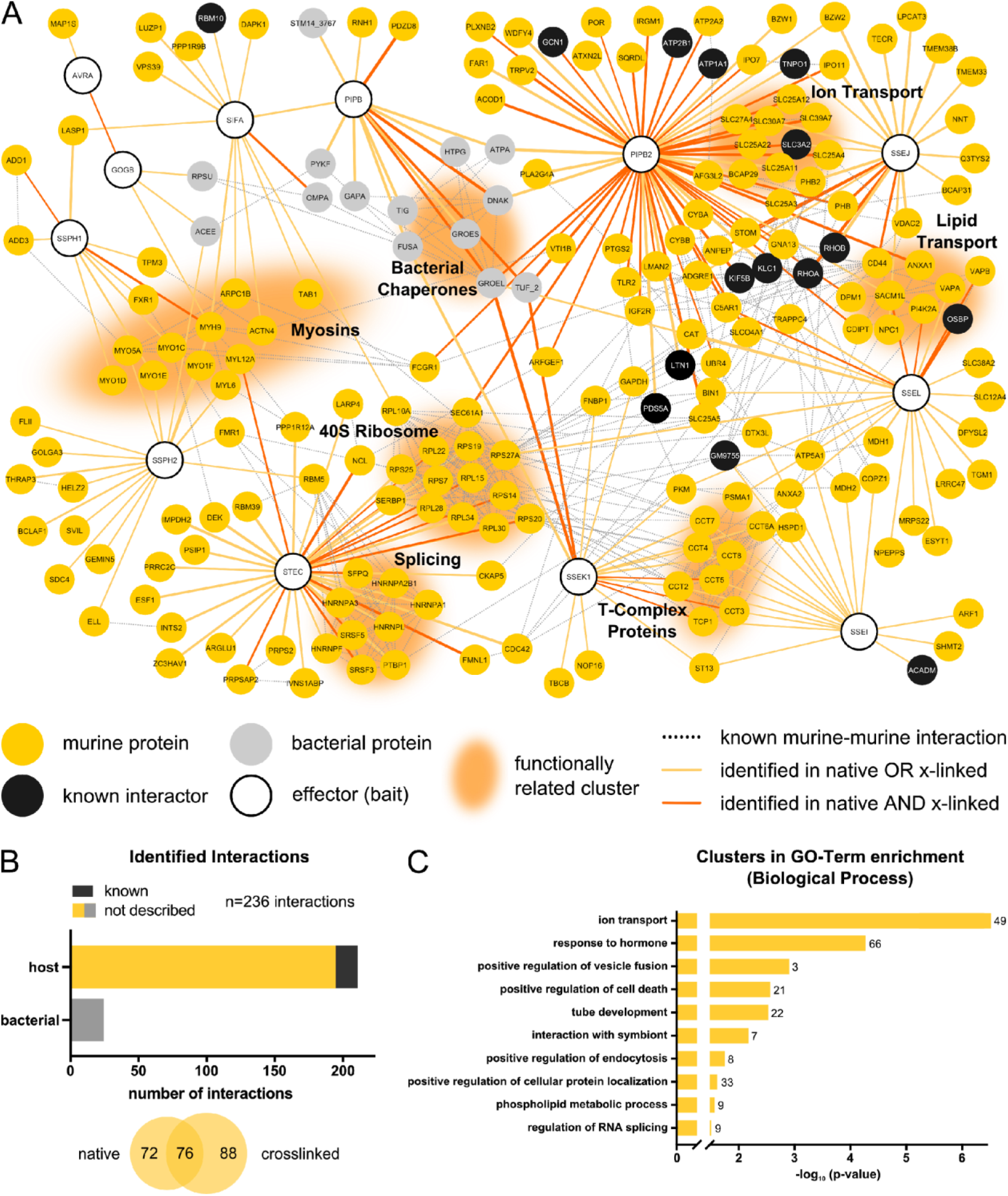
*S*Tm effector-host target physical interactions in RAW264.7 macrophages. (A) Network of PPIs identified between 12 *S*Tm effectors and their target proteins in RAW264.7 cells at 20 hpi. Note, only effectors that were identified as bait in AP-QMS and that had target proteins passing the criteria described in Figure 1B are depicted in this network. Host proteins from RAW264.7 cells are shown in gold (interaction not yet described) or black (previously identified interactions; see Table S4). *S*Tm proteins that were identified in pulldowns are depicted in grey. The color of the edge between two nodes denotes the conditions interaction captured, the edge thickness is proportional to the fold change (Log_2_). Functionally related clusters are grouped and annotated accordingly. The network was generated using Cytoscape version 3.7.2 (Shannon et al. 2003). Murine-murine, as well as bacterial-bacterial functional interactions were extracted from the built-in STRING DB version 11 (Szklarczyk et al. 2019) protein query for *Mus musculus* and *Salmonella* with a confidence cutoff of 0.7. (B) Overview of identified PPIs in RAW264.7 cells at 20 hpi. Hits are grouped according to whether they are of murine or *S*Tm origin (upper histogram), or according to whether they were detected in native or cross-linked pulldown samples (lower Venn diagram). (C) GO-term analysis for biological processes which are enriched among all identified PPI partners. GO-term clusters are ordered according to the significance of their enrichment (negative logarithmic, Benjamini-Hochberg corrected for multiple testing) (Benjamini and Hochberg 1995; Bindea et al. 2009) and top 10 GO-clusters are displayed. n signifies the number of proteins present in the respective cluster. Enrichments were normalized to the combined background proteome from AP-QMS experiments. A full list of identified enrichments can be found in Table S6.

**Figure 3.**
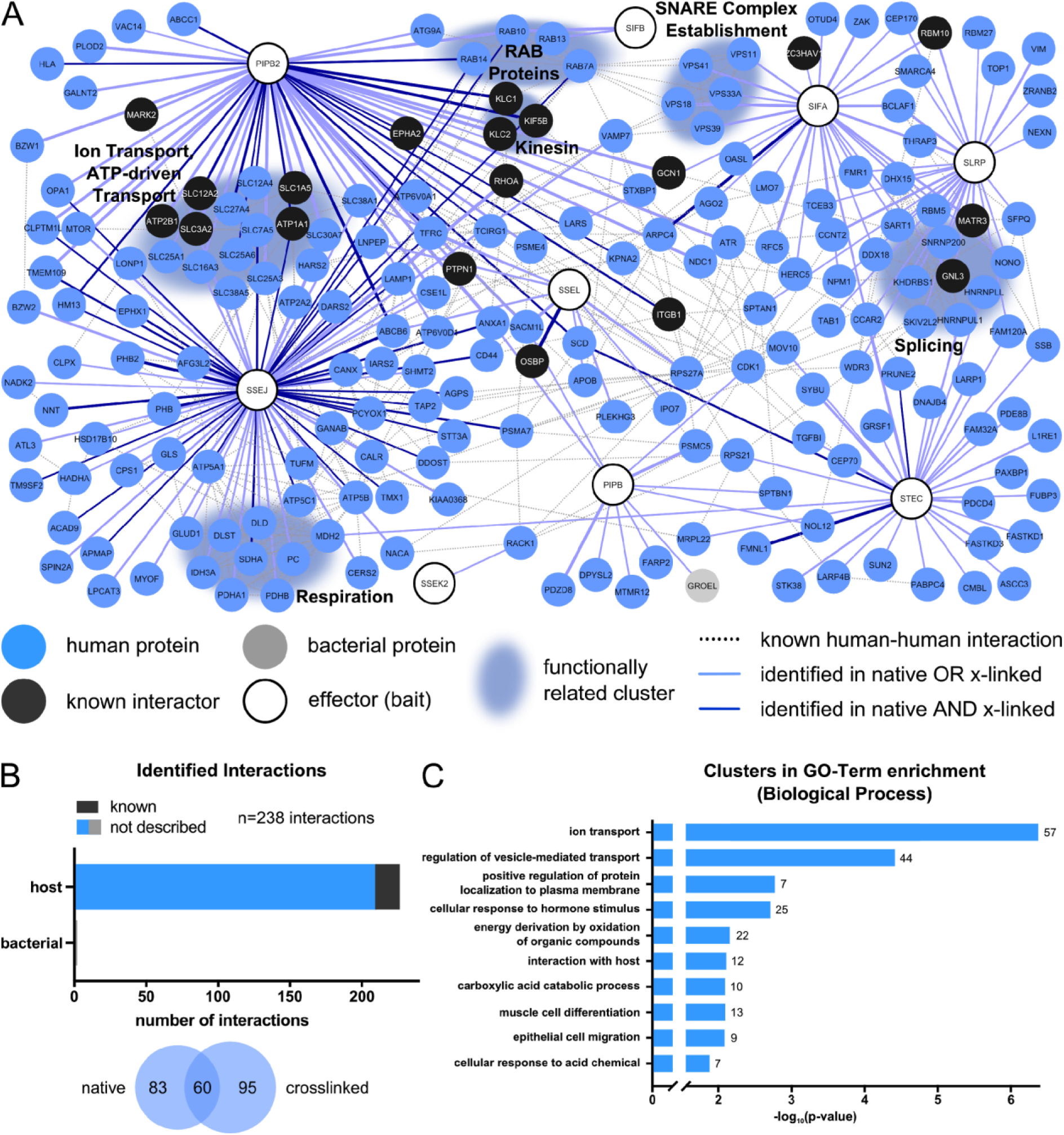
*S*Tm effector-host target physical interactions in HeLa cells. (A) Network of PPIs identified between 9 STm effectors and their target proteins in HeLa cells at 20 hpi. The same requirements and thresholds as described in Figure 2A were applied to select the nodes in the network. Host proteins from HeLa cells are displayed in blue (interaction not yet described) or black (previously identified interaction). *S*Tm proteins that were identified in pulldowns are depicted in grey. Edge formatting and network generation were performed as described for Figure 2A, with the exception that human data from STRING were used for generating functional interactions between host proteins. (B) Overview of identified PPIs in HeLa cells at 20 hpi - as in Figure 2B. (C) GO-term analysis for biological processes which are enriched among all identified interaction partners – performed and shown as in Figure 2C.

Rather than effectors interacting exclusively with a single host protein, we detected several effectors co-purifying with many host targets, such as PipB2, which had 59 in RAW264.7 cells and 48 PPIs in HeLa cells (Figure S7A). This implies that pleiotropic effectors may be the norm in bacterial pathogens, rather than the exception (Takahashi-Kanemitsu, Knight, and Hatakeyama 2020; Hamon et al. 2012). For example, SteC, a well-known *S*Tm effector with kinase activity, has been previously implicated in actin remodeling around the SCV (Poh et al. 2008). Here, SteC displayed several PPIs with host proteins related to mRNA splicing in both cell types, suggesting a potential, additional regulatory role in host-transcript splicing.

To check whether effectors target specific biological processes, in addition to overlaying human or murine functional interactions in the networks (Figures 2A and 3A), we performed GO-term enrichment on their targets (Figures 2C and 3C, Table S6). Ion transport and vesicle-mediated transport or fusion were among the most enriched targets in both cell lines (Figures 2C, 3C and S7B). The former stemmed mainly from interactions of PipB2 and SseJ with Small Solute Carrier proteins and ATP-dependent transporters, and the latter from interactions with many *S*Tm effectors. Other processes were specific to the cell line. Cytoskeleton-dependent transport, occurring mostly through the interaction of SspH1 and SspH2 with myosins, and lipid transport were specifically enriched in macrophages (Figure 2A and 2C, Table S6). Both processes were previously described to play important roles in SCV maintenance (Wasylnka et al. 2008; Nawabi, Catron, and Haldar 2008; Arena et al. 2011). In contrast, oxidation of organic compounds and respiration were prominent in epithelial cells, mainly due to interactions of SseJ, as well as specific interactions to RAB proteins (SifB, SseJ and PipB2) and to the SNARE complex (SifA; Figures 3A and C). These host machineries have been implicated in *S*Tm infection before (Stévenin et al. 2019; Kyei et al. 2006; Rzomp et al. 2003; Stein, Müller, and Wandinger-Ness 2012), however physical interactions *via* these specific effectors were not previously reported.

A notable feature of both PPI networks was that several effectors converged on the same host protein complexes/processes with myosins, ion transport, cholesterol transport, 40S ribosome and the T-complex being the most prominent hubs targeted by more than one effector (Figures 2A and 3A). In some cases, multiple effectors targeted the exact same host protein, such as myosin MYH9, which was bound by SspH1, SspH2, GogB and SifA in RAW264.7 cells (Figure 2A). This highlights the potential for effector co-operation on the same host cellular process (Figure S7C), which may occur simultaneously or in a parallel fashion. Interestingly, we also observed a number of effector-effector interactions (GogB-AvrA, PipB-SifA). Although some may be indirect and mediated through common host targets, this reinforces the notion that effectors converge on the same host processes and work cooperatively to hijack them. For example, both AvrA and GogB are known to impose an anti-inflammatory effect on host cells during *S*Tm infection. AvrA dampens JNK signaling via MKK7 (Du and Galán 2009), thereby reducing apoptosis (Jones et al. 2008), whereas GogB acts on NFkB by inhibiting degradation of IFkB (Pilar et al. 2012). Even though no common target for these two effectors has been described, the finding that they physically interact indicates a direct collaboration of AvrA and GogB in the regulation of inflammation.

One advantage of systematic studies is that common contaminants of pull-downs can be identified and normalized out during data analysis (see Experimental Procedures). This allows identification of specific interactions with targets that would normally be disregarded. For example, we detected 25 effector-bacterial PPIs in macrophage cells, e.g. PipB-DnaK, PipB-GroEL, PipB-STM14_3767 (Figure 2B). In order to exclude the possibility that these PPIs are due to partial bacterial lysis during infection or harvesting, which results in bacterial cytoplasmic proteins contaminating the host cytoplasmic fraction, we validated the presence of GroEL in the host cytoplasm during infection using a GroEL polyclonal antibody. Consistent with previous reports showing GroEL is secreted by *Bacillus subtilis, Helicobacter pylori* and *Francisella novicida* (Yang et al. 2011; González-López et al. 2013; Pierson et al. 2011; *McCaig, Koller, and Thanassi 2013), we detected GroEL* within the host lysate (Figure S8). This cannot simply be explained by bacterial lysis, as another abundant bacterial protein, RecA, was only detected in the bacterial cell pellet. This suggests that GroEL is secreted into host cells during infection and could play a role in effector functionality in the host cytoplasm. We obtained similar results for STM14_3767, a putative acetyl CoA hydrolase (Figure S8).

In summary, we recovered both previously identified PPIs and a plethora of new ones. Most STm effectors have multiple host targets, but in general, effectors converge to target the same processes in the host. Based on common host targets, we were able to draw new associations between specific effectors, which we anticipate will promote a deeper understanding of the complex interplay between effectors during infection.

### Strong interactions can be validated by reciprocal pull downs on the host target

The majority of PPIs we identified were cell line-specific (418/446, Figure 4A), which prompted us to investigate the underlying reasons for such differences (Figure S9, Table S7). About one third of the PPIs that were detected specifically in one cell type were due to the lack of detectable expression of that protein in the other cell line (Figure S9H, I, K and L). However, most cell-type specific PPIs had similar abundance in both cell lines (Figure S9G,J). The remaining differences can be due to false negatives and/or reflect differences in infection cycle in epithelial cells and macrophages – *S*Tm can escape and proliferate in the cytoplasm of epithelial cells, but not of macrophages (Knodler et al. 2010; Castanheira and García-Del Portillo 2017). Taken together, these results indicate that effector-host PPIs are largely cell-type specific, and only partially due to differences in protein expression.

**Figure 4.**
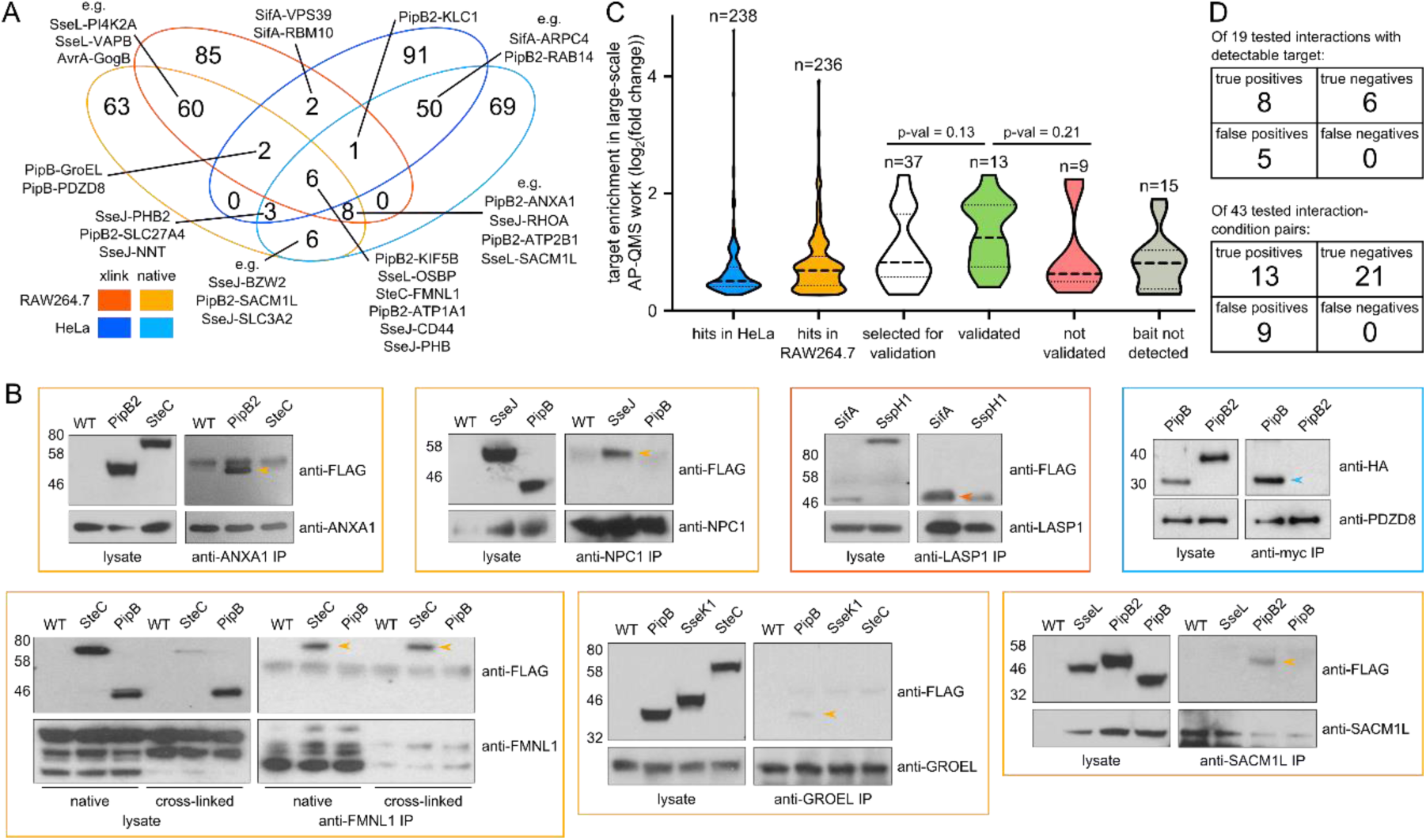
Comparison of *Salmonella* interactomes in RAW264.7 and HeLa cells, and reciprocal PPI validation. (A) Venn diagram comparison of PPIs across the two cell lines and conditions. (B) Reciprocal pulldowns using antibodies specific to host targets were used to validate PPIs detected in the AP-QMS screen. Antibodies to host proteins were added to Triton-X100 (0.1%) solubilized and centrifuged host cell lysates infected with indicated tagged effector strains for 20 h. Antibodies were then bound to Protein A/G beads, washed and eluted by boiling in Laemmli buffer. Effectors with similar expression levels were used in parallel pulldowns as negative controls. The PipB-PDZD8 reciprocal pulldown was performed by infection of HeLa cells expressing transfected myc-tagged PDZD8 with *S*Tm Δ*pipB* cells expressing PipB-2HA *in trans*; Δ*pipB2* expressing PipB2-2HA *in trans* was used as a negative control. For each reciprocal pulldown, two independent experiments were performed, except for LASP1 and GroEL pulldowns, which were performed once. Pulldown results were visualized by western blots with an antibody against the epitope tag fused to STm effector (anti-FLAG or anti-HA). One exemplary blot per interaction is shown, all blots and raw images are located in the Supplementary Material. Colored box around the Western Blot image indicates the cell background and condition tested. Validated interactions are indicated by arrows. (C) Violin plots of log_2_ fold enrichments in AP-QMS for all effector-target proteins selected to be tested by reciprocal pulldowns using the host protein as bait (white), those that could (green) or could not (red) be validated, and those where the bait was not detected in the reciprocal pulldown (grey). Dotted lines indicate median (bold) and interquartile range (light). For significance testing, two-sided T-test with Welsh correction was used, p-values are indicated. For comparison, the enrichments of all interactions identified in HeLa cells (blue) and RAW264.7 (orange) are shown. All tested interactions, their fold enrichments, as well as the respective results of reciprocal pulldowns are summarized in Table S8. (D) Tables summarizing the validation outcome with respect to the total number of assessed interactions. In the first table, interactions are considered irrespective of condition or cell line, i.e. an interaction is validated if it can be reproduced in at least one condition/cell line, in the second case each cell line and condition is taken as separate experiment.

Several PPIs were specifically identified in the presence of the crosslinker (Figures 2A, 3A and 4A). For example, SifA interacted with VPS39 and RBM10 only after crosslinking in both cell types, suggesting that these interactions may be transient. The only partial, though highly significant overlap (p-value < 0.0001, Fisher’s exact test) between native and crosslinked data could have additional reasons: a) loss in the efficiency of bait pulldown after crosslinking; b) increased background/poorer signal-to-noise in crosslinking experiment (Figure S2, S9C and F); c) differences in sample preparation and increased incubation times impacting the recovery of PPIs; and d) false negatives due to stringent thresholds, although our analysis tried to rectify this. A number of interactions were conserved across backgrounds and pulldown conditions, indicating strong interactions. We suspect that PPIs found in at least three of the four conditions indicate false negatives in the fourth condition. Among the conserved interactions, several were novel, e.g. SteC-FMNL1, PipB2-ATP1A1, PipB2-ANXA1, SseJ-CD44, SseL-SACM1L or PipB-GroEL.

To assess the validity of our newly identified interactions, we selected a subset of 12 host targets, which amounted to 22 distinct effector-host protein interactions – 37 PPIs taking into account all different conditions (native vs. cross-linked, cell line) – and sought to validate their interactions with the respective *Salmonella* effectors reciprocally (Table S8). The host targets were selected to span both weak and strong enrichment scores, as well as varying degrees of conservation of interactions throughout the different conditions tested (Figure 4B and C, Table S8). To test for reciprocal interactions, we pulled down on the host protein during *S*Tm infection using specific antibodies (see Experimental Procedures). In total, we could successfully pulldown 7 out of the 12 host target proteins, covering 13 of the 22 distinct PPIs (or 22 of the 37 tested conditions). In these cases, we could successfully recapitulate the orthogonal pulldown of the *S*Tm effector for 8 out of the 13 possible PPIs (61.5%) in at least one condition (13 of all the possible 22 conditions, i.e. 59.1% could be validated). We used a non-cognate *S*Tm effector of similar translocation level as a negative control (Figure 4B-D, summarized in Table S8). Note that these pulldowns were performed in a cell population containing 20-40% infected cells. Furthermore, even in infected cells the protein levels of translocated *S*Tm effectors are much lower than that of host proteins (Selkrig et al. 2018). Consequently, the majority of the target protein is unbound by the *S*Tm effector, either because the target protein comes from uninfected cells or because it is in large excess over the effector. Consistent with such an increased difficulty in capturing effector-host protein interactions by pulling down on host proteins, we observed that stronger PPIs (higher fold changes in screen) were more readily verifiable via reciprocal pulldowns (Figure 4C). In summary, we could recapitulate most of the newly identified effector-host protein interactions we tested using an orthogonal, but less sensitive approach. This suggests that many of the interactions we report here are also occurring during infection.

### SseJ and SseL cooperate to regulate intracellular cholesterol trafficking via NPC1

From our AP-QMS analysis, we identified “phospholipid metabolic process” and “positive regulation of vesicle fusion” as enriched GO-terms in RAW264.7 macrophages, and “regulation of vesicle-mediated transport” in HeLa cells, all of which comprise proteins involved in lipid and more specifically, in cholesterol trafficking. Host proteins required for cholesterol trafficking including OSBP, NPC1, VAPA/B, SACM1L were associated with multiple effectors. These interactions were predominantly mediated by the effectors SseJ, SseL and PipB2 in both HeLa cells (Figure 3A and S7B) and RAW264.7 macrophages (Figure 2A and S7B). As SseJ esterifies cholesterol (Nawabi, Catron, and Haldar 2008), we probed more carefully its connection with cholesterol transport by performing AP-QMS after crosslinking in HeLa cells in triplicate, and analyzed the samples with the corresponding untagged controls in the same TMT run (all results are summarized in Table S9). Combining all replicates into a single TMT run increases the sensitivity in detecting low abundant PPIs (because sample complexity is greatly reduced compared to multiplexing with 9 other pulldowns). This enabled us to detect a PPI between SseJ and the effector SseL (Figure 5A), which is in line with recent evidence demonstrating functional cooperation between these two effectors to promote SCV stability via interactions with OSBP (Kolodziejek et al. 2019), a lipid transfer protein that controls cholesterol/PI4P exchange between the ER and Golgi (Mesmin et al. 2017). Consistently, OSBP co-purified with SseJ and SseL in both cell lines (Figures 2A, 3A and 5A).

**Figure 5.**
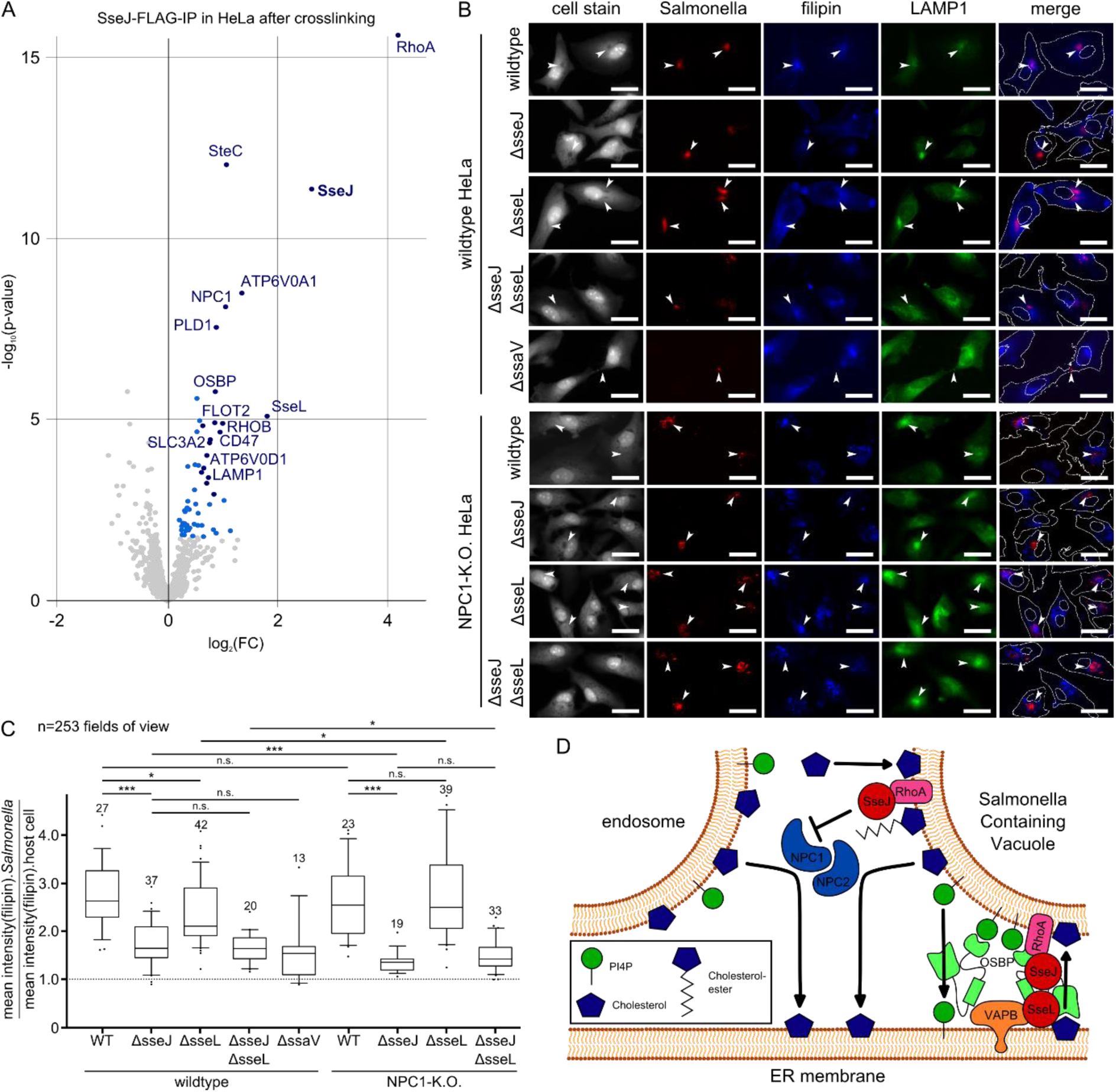
SseJ and SseL influence intracellular cholesterol trafficking. (A) Volcano plot showing enrichments after crosslinked pulldown of STF-tagged SseJ in HeLa cells at 20 hpi compared to untagged control (wildtype infection). Three replicates for SseJ-STF and wildtype were measured in a single TMT run. Dark blue: FC > 1.5, p-value < 0.001; light blue: FC > 1.2, p-value < 0.01. Apart from the bait (SseJ), previously described interaction partners RhoA and OSBP, a number of new host targets, and the *S*Tm effectors SseL and SteC are significantly enriched. All hits are summarized in Table S9. (B) Representative microscopy images (20x magnification) at 12 hpi with mCherry-expressing *S*Tm strains in HeLa cells (wildtype and NPC1-knockout). Draq5 staining is displayed in grey, *S*Tm in red, filipin (stains unesterified cholesterol) in blue and LAMP1 in green. Arrows indicate the location of *S*Tm microcolonies across the fluorescence channels. The last column displays a merge of mCherry and filipin signals, with the outlines of the cell periphery and the nuclei drawn in white. Scale bar: 30 µm. (C) Quantification of B. A total of 253 manually inspected fields of view across two independent experiments with four technical replicates in each run, were analyzed. For quantification, the average filipin intensity in regions of co-localization of intracellular STm with LAMP1 staining (to exclude cytosolic bacteria) was divided by the average filipin intensity measured within the cell mask. The analysis was performed by field view (n shown in boxplots). Field of views contained on average 20 infected cells. Boxplots (median and interquartile range) with whiskers spanning Q10 to Q90 are displayed. For statistical analysis, T-test with Welsh correction was used and significance indicated as follows ***: p-value < 0.001, *: p-value < 0.05, n.s.: not significant (p-value > 0.05). (D) Model of the interdependence of SseJ, SseL and cellular targets in cholesterol trafficking during infection. NPC1 and NPC2 are involved in recycling cholesterol from endosomes, as well as presumably from the SCV, thereby replenishing the pool of cholesterol in other compartments, such as the ER. The presence of cholesterol in the ER membrane provides a substrate pool for OSBP, which transports cholesterol to the TGN (and presumably to the SCV) in exchange for PI4P. This directional cholesterol trafficking is presumably enhanced mainly by SseJ, and to a lesser extent by SseL. In addition to its binding to OSBP, SseJ, which localizes to the SCV in a RhoA-dependent manner, has been shown to bind cholesterol independently of other factors (Nawabi, Catron, and Haldar 2008), which is in line with our finding that SseJ is the primary effector in cholesterol recruitment to the SCV. The role of SseL in enhancing cholesterol trafficking to the SCV is NPC1-dependent, but does not necessarily rely on direct PPI.

In addition to detecting the recently reported SseJ-OSBP interaction (Kolodziejek et al. 2019), we observed an interaction between SseJ and the Niemann-Pick disease type C1 protein (NPC1) (Figure 5A). NPC1 plays a critical role in cholesterol trafficking (Pfeffer 2019). We thus wondered whether SseJ and SseL alter cholesterol trafficking via NPC1. To probe this and validate the roles of SseJ and SseL in this process, we infected HeLa cells with wildtype and mutant *S*Tm and stained with filipin at 12 hpi. Filipin stains unesterified cholesterol and is commonly used to assess the intracellular distribution of cholesterol (Maxfield and Wüstner 2012; Wilhelm et al. 2019). Cholesterol was recruited to the SCV upon infection with wildtype *S*Tm (Figure 5B and C). We assessed co-localization between cholesterol and the SCV by calculating the ratio between filipin signal at the site of the SCV and the overall filipin signal per cell. This means that a random cholesterol distribution throughout the cell results in a ratio of 1, stronger co-localization in values >1 and exclusion of filipin at the SCV in values <1. This ratio was reduced strongly upon infection with an Δ*sseJ* mutant (wildtype median = 2.62 and Δ*sseJ* = 1.64). Infection with Δ*sseL* bacteria also reduced cholesterol accumulation at the SCV, albeit to a lesser extent (median = 2.11). Interestingly, the double Δ*sseJ*Δ*sseL* mutant and the SPI-2 secretion system null mutant (Δ*ssaV*) resulted in low SCV cholesterol accumulation comparable to Δ*sseJ* bacteria (Figure 5C), suggesting cholesterol accumulation at the SCV is largely driven by SseJ.

In order to explore the role of NPC1 in this process, we infected NPC1 KO cells with wildtype *S*Tm. This resulted in cholesterol accumulation at the SCV comparable to that observed in wildtype HeLa cells, despite NPC1 KO cells exhibiting pronounced endosomal cholesterol accumulation, as previously reported (Tharkeshwar et al. 2017). Wildtype *S*Tm was able to overcome this aberrant endosomal cholesterol localization in NPC1 KO cells and accumulated cholesterol at the SCV at similar levels to those detected in wildtype HeLa cells. Interestingly, the Δ*sseL* mutant no longer conferred reduced cholesterol accumulation compared to wildtype HeLa cells. This suggests that the minor role of SseL in SCV cholesterol accumulation operates *via* NPC1. Infection with Δ*sseJ* or Δ*sseJ*Δ*sseL* mutants further aggravated the absence of cholesterol from the SCV relative to that observed in wildtype HeLa cells (Figure 5C). Taken together, these findings suggest a complex interplay between SseJ and SseL, and multiple host target proteins (e.g. NPC1, OSBP) to modulate cholesterol trafficking during infection. While the mild impact of SseL-mediated recruitment of cholesterol to the SCV requires NPC1, this is likely not caused by direct physical interaction, but rather through a functional dependence and indirect interactions with OSBP and SseJ (Figure 5D).

### PipB interacts with PDZD8 and recruits it to the SCV

We identified a strong interaction between PipB and the PDZ-domain containing protein 8 (PDZD8) in both HeLa and RAW264.7 cells (Figure 2A, 3A and 4A). PDZD8 is a paralog of the ERMES (ER-mitochondria encounter structure) component Mmm1 and was recently shown to play a functional role at ER-mitochondrial contact sites by regulating Ca^2+^ dynamics in neurons (Wideman et al. 2018; Hirabayashi et al. 2017). We were able to verify this interaction *via* ectopic expression of EGFP-tagged PipB in HeLa cells and MS identification of PDZD8. PDZD8 did not co-IP with EGFP-PipB2, its effector ortholog (Figure S10), thus demonstrating the specificity of the PipB-PDZD8 interaction.

We then sought to map the PipB-PDZD8 PPI and its cellular localization during infection. Ectopic expression of EGFP-PipB resulted in co-localization with PDZD8 at the ER, based on the ER resident marker protein disulphide-isomerase (PDI) (Figure 6A). To examine the PDZD8-PipB interaction in an infection context and after PipB translocation, we infected PDZD8-myc expressing HeLa cells with *S*Tm Δ*pipB* bacteria expressing PipB-2HA *in trans*. We observed a striking accumulation of PDZD8 specifically at the SCV, but not on *Salmonella*-Induced Filaments (SIFs), based on its partial overlap with PipB-2HA and the SCV/Sif marker protein, LAMP2 (Figure 6B). Recruitment of PDZD8 to the SCV was PipB-specific, as PDZD8-myc was not recruited to the SCV upon infection of HeLa cells with Δ*pipB2* pPipB2-2HA bacteria (Figure 6C). These findings demonstrate that PipB specifically recruits PDZD8 to the SCV during infection (see also Figure S10).

**Figure 6.**
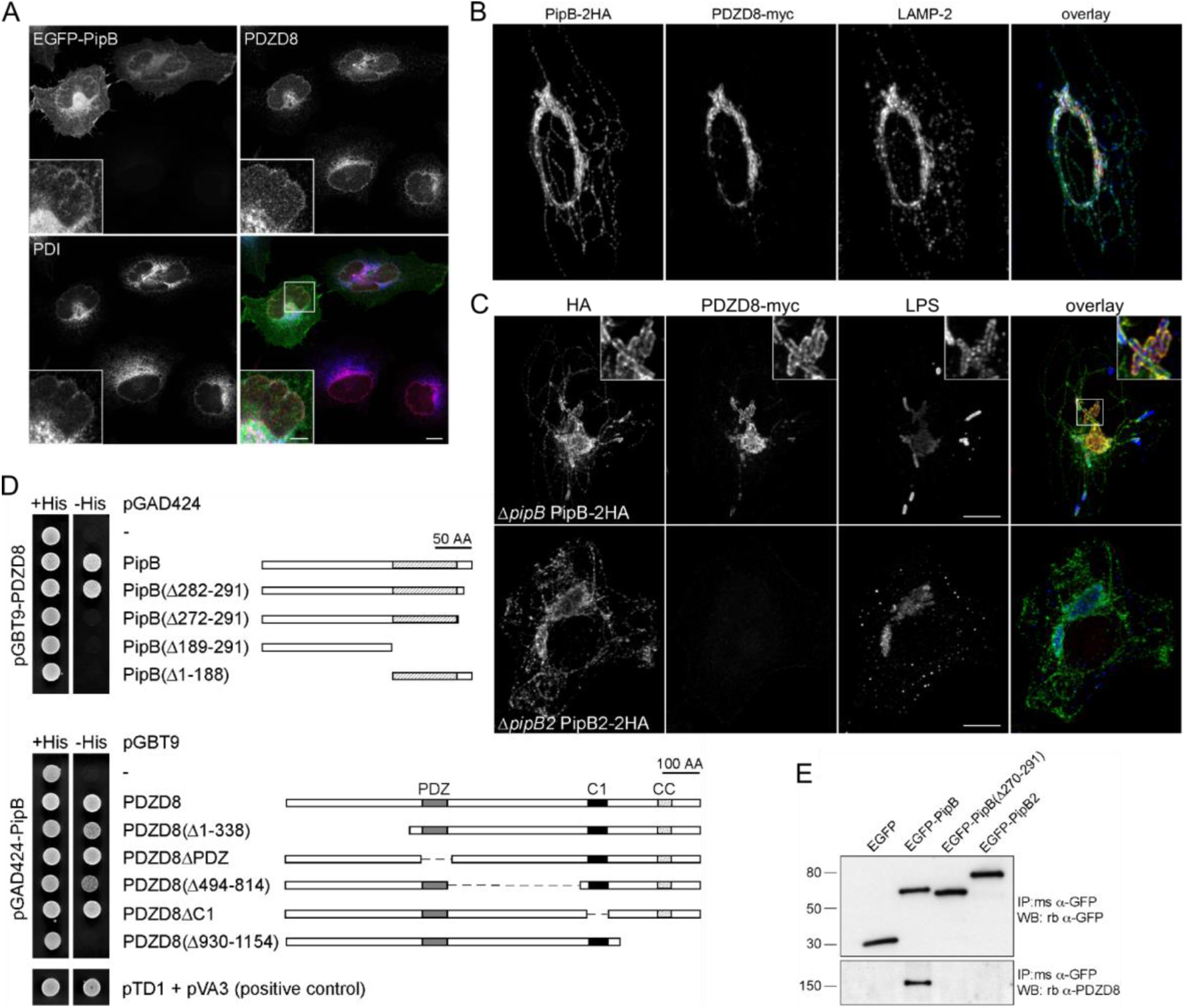
PipB recruits PDZD8 to the SCV during infection. (A) Immunofluorescence microscopy of EGFP-PipB, endogenous PDZD8 (stained with antibody) and Protein Disulphide-Isomerase (PDI) shows co-localization of these proteins at the ER after transfection of HeLa cells. Main scale bar: 5 µm, and the inset: 2 µm. (B) Fluorescence microscopy image showing that HA-tagged PipB localizes to the SCV and SIFs, as shown by staining for LAMP-2. HeLa cells were transfected with PDZD8-myc and infected with STm Δ*pipB*, carrying a plasmid expressing PipB-2HA, and imaged at 12 hpi. PDZD8 co-localizes at the SCV surface, yet not along the SIFs. Scale bar: 5 µm. (C) Fluorescence microscopy of HA-tagged PipB and PipB2 with myc-tagged PDZD8 and LPS to stain *Salmonella*. HeLa cells transfected with PDZD8-myc and infected with STm Δ*pipB* pPipB-2HA or Δ*pipB2* pPipB2-2HA, 12 hpi. Localization of PDZD8 to the SCV is dependent on PipB expression *in trans*, but not PipB2. Scale bar: 5 µm. (D) Yeast two hybrid assay with truncated versions of PipB or PDZD8. Direct interaction between the two proteins, as indicated by growth in -His conditions, is abolished by deletion of the 20 amino acid C-terminus of PipB. Numbers indicate the deleted residues. In PDZD8, deletion of the PDZ- or C1-domains does not impair interaction with PipB, but deletion of the 225 C-terminal amino acids does. (E) Western Blot after immunoprecipitation from HeLa cells transfected with EGFP, EGFP-PipB, EGFP-PipB(Δ270-291) and EGFP-PipB2 fusions. Anti-GFP immunoprecipitation was analyzed by immunoblotting for endogenous PDZD8 using anti-PDZD8 peptide antibodies and anti-GFP antibodies. The PipB-PDZD8 interaction requires the last 20 amino acids of PipB. PipB2 was used as negative control to test the PipB-PDZD8 interaction specificity.

In order to map the PDZD8-PipB interaction in further detail, we created a series of PipB and PDZD8 truncations and tested their ability to interact by yeast two hybrid (Y2H). For PipB, truncating the C-terminal 20 amino acids (Δ272-291) resulted in disruption of PipB-PDZD8 binding (Figure 6D). The last 20 amino acids alone, however, were not sufficient for the interaction with PDZD8, as deletion of the N-terminal 188 amino acids (Δ1-188) also disrupted PDZD8 binding. As for PDZD8, a critical segment within its C-terminal 224 amino acids (Δ930-1154) that contains a predicted coiled-coil domain was required for the interaction with PipB (Figure 6D). We verified the importance of the C-terminus of PipB in mediating the interaction with PDZD8 by transfecting HeLa cells with EGFP-effector fusions. Consistent with the Y2H data, endogenous PDZD8 co-immunoprecipitated with PipB and not PipB2, whereas deletion of the C-terminal 22 amino acids of PipB abolished its interaction with PDZD8 (Figure 6E). These results illustrate an important role for the C-terminal domains of PipB and PDZD8 in mediating their physical interaction. This concurs with the previous observation that the functional divergence of PipB and PipB2 is due to sequence divergence in their extreme C-termini (Knodler and Steele-Mortimer 2005).

### SteC targets FMNL1 to promote actin polymerization

We identified a novel PPI between the effector kinase SteC and a formin-line protein, FMNL1, which is highly expressed in macrophages (Yayoshi-Yamamoto, Taniuchi, and Watanabe 2000). In all conditions and cell lines tested, FMNL1 co-purified with SteC and this PPI was also verified by reciprocal pulldowns in both HeLa and RAW264.7 cells (Figure 4). To examine whether the SteC-FMNL1 PPI is the result of a direct PPI between these two proteins, we purified full-length SteC, a catalytic inactive mutant SteC_K256H_ and the N-terminal domain of FMNL1_1-385_, and tested for complex formation by size-exclusion chromatography. SteC and FMNL1 alone migrated as multimeric species (Figure 7A; blue and orange traces, respectively), but when pre-incubated together, a portion of FMNL1_1-385_ co-migrated with SteC forming a higher molecular weight complex (Figure 7A; green trace). This was also true for SteC_K256H_. Thus, SteC directly binds to the N-terminus of FMNL1 independent of its kinase activity.

**Figure 7.**
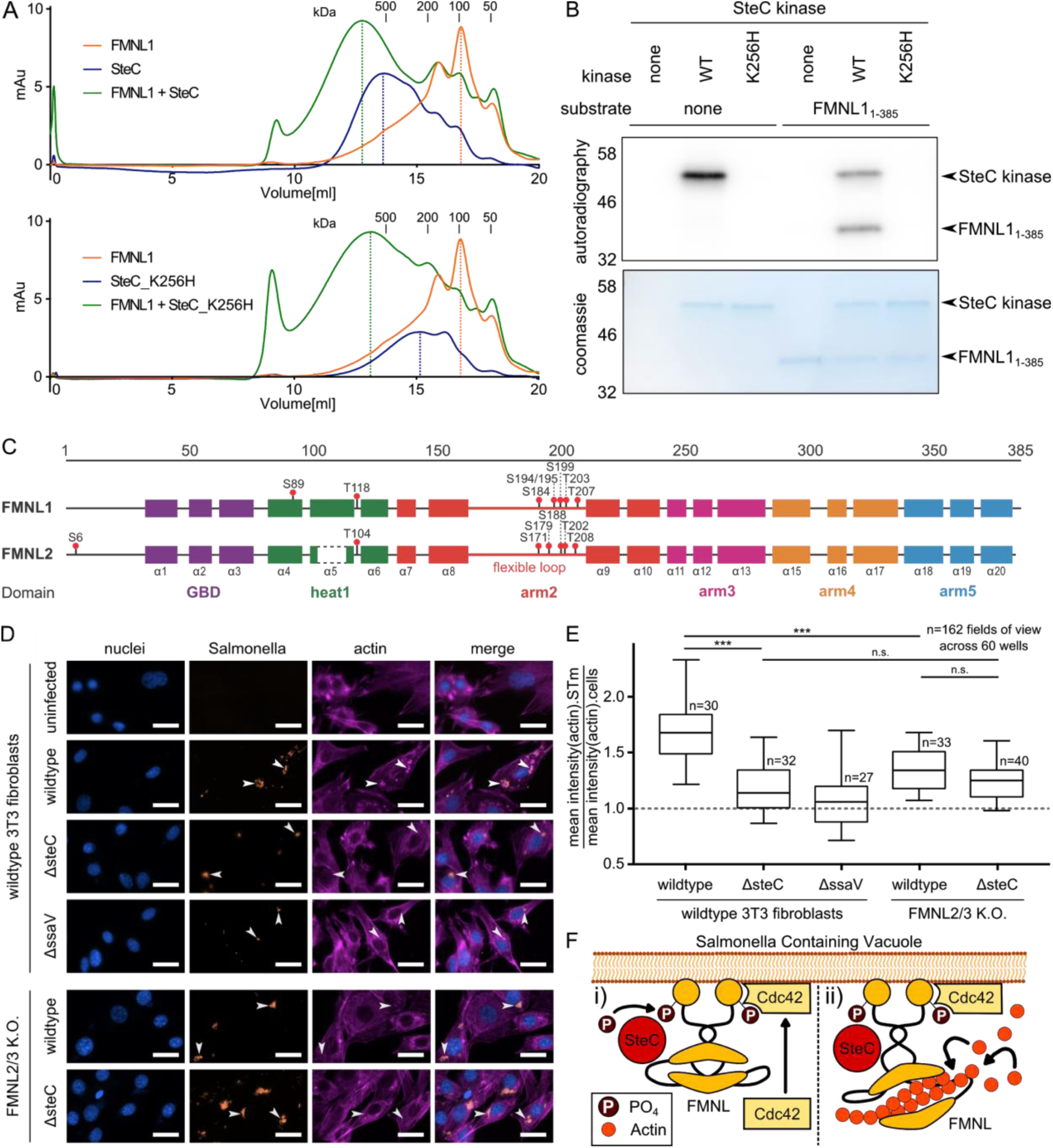
SteC directly targets FMNL proteins to promote actin cytoskeleton rearrangements. (A) Size exclusion chromatograms obtained from purified recombinant FMNL1_1-385_ (orange traces), SteC (blue trace, upper panel) or catalytically inactive SteC_K256H_ (blue trace, lower panel). Pre-incubation of FMNL1_1-385_ with SteC (green trace, upper) or SteC_K256H_ (green trace, lower). A shift in elution volume in the mixed sample compared to the individual purified proteins, as indicated by the dotted lines demonstrates direct interaction of both SteC and SteC_K256H_ to FMNL1. Retention times corresponding to specific molecular weights were determined using Bio-rad protein standard (1.35 - 670 kDa). (B) Autoradiography after *in vitro* kinase assay. FMNL1_1-385_ was purified and incubated with purified SteC kinase, as well as SteC_K256H_ in the presence of radioactively labelled [^32^P]-γ-ATP. Samples were separated by SDS-PAGE, transferred to a PVDF membrane and imaged in a phosphoimager. Protein inputs were similar as shown by Coomassie blue staining. Only catalytic active SteC is capable of autophosphorylation, as well as phosphorylating FMNL1 *in vitro* (bands indicated by arrows). (C) Protein maps of FMNL1 and FMNL2, including functional regions and secondary structure elements, as well as phosphosites identified in the *in vitro* kinase assay, followed by phosphoproteomics. Comparison between the two maps shows that phosphorylation by SteC occurs mostly in the flexible loops of FMNL1 and FMNL2. Results are summarized in table S10 (also SteC autophosphorylation sites). (D) Representative fluorescence microscopy images after infection of 3T3 fibroblasts (8 hpi) with different mCherry-expressing STm strains. Images were obtained after staining with DAPI (blue) and phalloidin (purple). Data from three independent experiments for FMNL2/3 knockout cells (in all experiments, both clones, 9.10 and 46.20 described in (Kage, Steffen, et al. 2017) were used, each in 20 wells per experiment), and two independent experiments for wildtype 3T3 fibroblasts spanning 162 fields of view (20 infected cells on average per view). Representative images are shown, and corresponding quantification is displayed in E. Arrows indicate intracellular STm microcolonies, as well as their position in other fluorescence channels. Scale bar: 30 µm. (E) Quantification of average actin signal intensity at the site of STm microcolonies divided by overall average actin signal intensity as a measure of co-localization between actin and STm. A total of 162 fields of view across 60 wells were analyzed and are here displayed as boxplots. Boxplots are drawn as in Fig. 5c. For statistical analysis, T-test with Welsh correction was used and significance is indicated ***: p-value < 0.001, n.s.: p-value > 0.05. (F) Model of SteC-FMNL interaction and functional relationship: (i) SteC binds FMNL subfamily formins directly, and independently of its catalytic activity, and is necessary and sufficient for its phosphorylation. (ii) The interaction between phosphorylated FMNL formins and Cdc42 induces actin polymerization (Kühn et al. 2015), and explains the actin bundling phenotype (as observed in fluorescence microscopy of infected 3T3 fibroblasts).

We then asked whether FMNL1 is a direct substrate of SteC. To test this, we performed an *in vitro* kinase assay in the presence of [^32^P]-γ-ATP. Consistent with previous reports, SteC was capable of auto-phosphorylation (Poh et al. 2008). In addition, when combined with FMNL1_1-385_, SteC, but not the catalytically inactive SteC_K256H,_ phosphorylated FMNL1 (Figure 7B). To identify the specific FMNL residues that are phosphorylated by SteC, we performed phosphoproteomics after an *in vitro* kinase assay for both FMNL1 and FMNL2, using SteC_K256H_ as negative control. Thereby we identified the SteC auto-phosphorylation sites and several phosphosites in similar domains of both FMNL proteins (Table S10), many located in the flexible loop of the armadillo repeat region (Figure 7C). Among other sites, S171 (FMNL2) and an equivalent site in FMNL1 (S184) were phosphorylated. This site has previously been shown to induce binding to Rho-family GTPase and FMNL regulator Cdc42 (Kühn et al. 2015).

SteC is required to induce actin bundling around the *S*Tm microcolony in 3T3 fibroblasts (Poh et al. 2008; Odendall et al. 2012; Imami et al. 2013). We therefore postulated that FMNL proteins may be required for this phenotype as they are known to promote actin polymerization (Bai et al. 2011; Heimsath and Higgs 2012; Block et al. 2012). Since FMNL1 had been considered to be most abundant in hematopoietic cells and low in expression in 3T3 fibroblasts (Kage, Winterhoff, et al. 2017), cells disrupted in the more ubiquitous FMNL2 and FMNL3 were used. As shown before (Odendall et al. 2012), actin bundling around the SCV was strictly dependent on SteC in 3T3 fibroblasts (Figure 7D). Interestingly, actin bundling around *S*Tm microcolonies was diminished in FMNL2/3 knockout fibroblasts no matter whether we infected with *S*Tm wildtype or Δ*steC* (Figure 7E), suggesting that SteC acts *via* FMNLs. Despite both *S*Tm strains not being able to induce substantial actin bundling in the absence of FMNL2 and FMNL3, there was still some residual bundle formation by SteC. We therefore examined more closely the expression of FMNL subfamily proteins in 3T3 cells using a newly available FMNL1-specific antibody. While FMNL2 and FMNL3 were abundant in control and absent in FMNL2/3 knockout cell lines, as expected, we could also detect a high molecular weight variant of FMNL1 in both cell lines (Figure S11). We suspect that the residual SteC-dependent actin bundling observed in FMNL2/3 knockouts can be ascribed to this FMNL1 expression. Taken together, these results demonstrate that SteC directly interacts with and phosphorylates FMNL subfamily formins. This could result in FMNLs binding to Cdc42 and inducing actin bundling at sites of *S*Tm microcolony formation (Figure 7F; see also Discussion).

## Discussion

In this work, we describe 421 novel PPIs, along with 25 previously described PPIs, between 15 different *S*Tm effectors after infection of two host cell lines (epithelial and macrophage). These interactions were identified during infection and physiological expression levels, using a quantitative proteomics-based approach (AP-QMS). We capitalized on the genetic tractability of *S*Tm and generated a library of 32 C-terminally tagged T3SS1 and T3SS2 effector strains (except SifA), which can be used in the future to probe different infection conditions (e.g. SPI-1 ON), cellular activation states (e.g. interferon-γ priming), spatiotemporal dynamics of effector localization and PPIs, and cell types (e.g. dendritic cells, intestinal epithelial cells). The majority of interactions we observed were cell type-specific (418 PPIs), with only 28 PPIs being conserved across the two cell-types. Although the stringency of our methodology may account for some of the differences, the differential expression levels of host targets (see Figure S9 and S11 for different FMNLs) and the different infection trajectories in epithelial cells and macrophages are more likely the reasons for this discrepancy. Most effectors co-purified with multiple host targets, several of which were related in function, uncovering an interconnected network of potentially overlapping functionalities between effectors. The functional relevance of this resource is exemplified by three vignettes of novel infection biology in cholesterol trafficking, organelle organization and actin rearrangements. The data provided can function as a rich resource for further investigating the complex interconnection between *S*Tm and host defense.

Several endeavors to map *S*Tm-host PPIs have been undertaken in the past (Auweter et al. 2011; Sontag et al. 2016). However, all were conducted outside the context of infection and typically after overexpression of an individual *S*Tm effector inside host cells. For example, one of the first systematic studies in the field, Auweter et al. ectopically expressed a panel of 13 effectors in HEK-293T cells and expressed and purified 11 effectors in *E. coli*. AP-MS in HEK-293T cells or HEK-cell lysates revealed 15 effector-target interactions, two of which (SseJ-RhoA and SseL-OSBP) (Auweter et al. 2011) were also identified in our study. In the report by Sontag et al., eight *S*Tm effectors were tested *in vitro* using AP-MS on purified effectors and RAW264.7 cell lysates (Sontag et al. 2016). Three of the effectors from this study are shared with our current study (SseL, SspH1 and SseI, also called SrfH), where for SseI, two interactions could be seen in both studies i.e. SseI-Gm9755 and SseI-ACADM. Interestingly, Sontag et al. identified various SLC proteins as targets of SseI and GtgA. We, however, detected several solute carrier proteins (SLCs) as targets of PipB2 and SseJ e.g. SLC25A5 and SLC25A11. It is plausible that SLCs are common targets of multiple *S*Tm effectors. SLCs have been linked to innate immunity, cytokine release, as well as bacterial and viral infections (Awomoyi 2007; Singh et al. 2016; Nguyen et al. 2018). It is thus conceivable that *S*Tm may modify SLC function to improve uptake into the host cell or to modulate inflammation. More recently, BioID was used to study effector-host interactions by tagging a panel of five effectors (PipB2, SseF, SseG, SifA, SopD2) with the biotin ligase BirA and overexpressing fusion proteins in HeLa cells by plasmid-based transfection (D’Costa et al. 2019). In the same study, the authors used AP-MS after ectopic effector expression. Although we tagged these same 5 effectors, due to limiting levels of translocated protein, we only assessed PipB2 and SifA by AP-QMS. Comparing the two studies, there is some overlap in host-protein targets: 4 proteins for SifA and 16 for PipB2, which are summarized alongside other previously described interactions in Table S4. The overexpressed effectors, the absence of an infection context and the stringent thresholds may account for the differences between the two studies. The authors did, however, find enriched processes (e.g. ion transport, SNARE complex, lipid metabolism, actin-related), which are congruent with our observations (Figure 2C, 3C and S7B), thus providing further validity of the importance of these processes in infection. Taken together, although there is some overlap between our study and past studies performed outside an infection context and for smaller sets of effectors, many of the PPIs are unique to our study and likely represent interactions that can only be captured in the context of infection.

Although our effector library contains a large number of known effectors, current effector knowledge may largely underestimate the full repertoire of proteins translocated or secreted by *S*Tm during infection (Li et al. 2018; Niemann et al. 2011). Recent proteomic methodologies have enabled unbiased profiling of secretomes of intracellular pathogens during infection (Mahdavi et al. 2014), and will be vital for uncovering the full repertoire of *S*Tm effectors translocated during infection. As more effectors are verified, they can be easily incorporated into this library and screened for PPIs.

Tagging can impede function or localization of some effectors, as previously shown for SifA (Brumell, Goosney, and Finlay 2002) – therefore, we adjusted the tagging strategy for SifA here. It will be important to assess whether the C-terminal modification introduced into these strains impacts effector translocation and function. We probed expression and translocation for all effectors, and we could detect 20 effectors in the host cytoplasm (28 were expressed). Although some of the remaining 12 may fail to translocate due to their C-terminal tag, we find it more likely that they are not translocated in sufficient amounts in the cell lines and/or time-points probed here. Introduction of a C-terminal tag may have led, in some cases, to poor stability/expression of otherwise abundant effectors, such as SseF and SseG. Of the 15 translocated effectors we could reproducibly detect by AP-QMS, there were 5 effectors for which we were unable to detect significantly enriched targets in at least one of the tested cell lines (GogB, SspH1, SspH2, SseK1 in HeLa cells and SlrP in RAW264.7 cells). In addition to tags compromising PPIs with host targets, other explanations could include promiscuous or transient interactions (many *S*Tm effectors are enzymes) or non-proteinaceous targets (lipid, DNA/RNA, metabolites) (Nawabi, Catron, and Haldar 2008; Knodler et al. 2009; McShan et al. 2016). In general, our inability to detect some PPIs rigorously described in literature, such as SifA-SKIP (Jackson et al. 2008; Diacovich et al. 2009; Zhao et al. 2015) or AvrA-MKK7 (Jones et al. 2008; Du and Galán 2009), could be due to tagging, conditions (20 hpi, stringent pull-downs) and cell lines used, MS-limitations (abundance or detection of prey) or false negatives of the method.

In order to capture transient PPIs, we used the crosslinker DSP, which resulted in both gains and losses of PPIs detected. In addition to transient interactions, these differences may be due to altered protein background, stringent thresholds for hit calling, competition for binding of targets, and differences in the experimental workflow. Past efforts to capture transient interactions have employed BirA-based approaches combined with formaldehyde crosslinking or AP-MS (stable complexes) combined with BioID (transient or proximal PPIs) to capture effector interactions (Mousnier et al. 2014; D’Costa et al. 2019). Combining such methods with our approach to map to host-pathogen PPIs during infection will likely add another layer of spatiotemporal complexity underlying host-pathogen PPIs. In addition, the use of catalytically inactive mutants of effector proteins may enhance the ability to capture target molecules. Combining pull-down approaches with lipidomics or metabolomics (Maeda et al. 2013; Saliba et al. 2014), may help to identify non-proteinaceous effector targets.

Several functionally related clusters were identified as targets of single or multiple effectors in our screen. One of these processes was cholesterol trafficking, which was mainly targeted by SseL, SseJ and PipB2 through interactions with a number of proteins involved in this process: OSBP, VAPA, SACM1L, PI4K2A, NPC1 and ANXA1. These links are in line with previous reports (Wyles, McMaster, and Ridgway 2002; Auweter et al. 2012; Mesmin et al. 2017; Kolodziejek et al. 2019). In this study, we provide supporting molecular evidence for the functional cooperation between the two effectors SseJ and SseL (Kolodziejek et al. 2019). Both are required to accumulate unesterified cholesterol at the SCV, which presumably makes it more stable (Kolodziejek et al. 2019). In NPC1 knockout cells however, the role of SseL, is fully mitigated, whereas the role of SseJ becomes dominant. These data suggest that SseJ and SseL cooperate to maintain cholesterol at the SCV through opposing effects that require NPC1. Interestingly, both SseJ and NPC1 localize to SIFs during infection (Ohlson et al. 2005; Drecktrah et al. 2008). It remains to be tested whether NPC1 recruitment to SIFs requires SseJ and/or SseL. Further work will also be required to elucidate the detailed molecular interactions that occur between SseL and SseJ and the proteins orchestrating cholesterol transport between organelles. Deletion of both *sseJ* and *sseL* has recently been shown to increase the fraction of cytoplasmic *S*Tm, indicating a role of these two effectors and the associated lipid trafficking in stabilization of the SCV (Kolodziejek et al. 2019).

We also identified a strong interaction between PipB and the host target PDZD8, a protein recently shown to be required for ER-mitochondrial and ER-lysosomal organelle contact sites in neurons, and for regulating Ca^2+^ levels therein (Hirabayashi et al. 2017). PDZD8 accumulates in contact sites between the ER and late endosomes/lysosomes in non-human primate kidney Cos-7 cells, together with Rab7 (Guillén-Samander, Bian, and De Camilli 2019). Interestingly, Rab7 was also enriched in PipB pulldowns, but remained just below our stringent significance thresholds (Table S2). We show that PipB binds to the C-terminal coiled-coil domain of PDZD8, which is the same region that mediates the Rab7-PDZD8 interaction (Guillén-Samander, Bian, and De Camilli 2019). In addition, PDZD8 has been identified as a moesin binding protein that impairs intracellular replication of Herpes Simplex Virus infection through regulation of the cytoskeleton (Henning et al. 2011). This connection to the cytoskeleton had also been described outside of the infection context (Bai et al. 2011). It is tempting to speculate that PipB promotes organelle tethering between the SCV, late endosomes/lysosome and the ER through interactions with Rab7 and PDZD8.

One of the strongest and most abundant interactions we detected was that between SteC and FMNL1. The kinase SteC had been linked to actin rearrangements during infection by modulation of MAPK signaling and HSP70 (Odendall et al. 2012; Imami et al. 2013). Yet, the effect attributed to SteC exceeded these interaction partners, indicating a missing piece in the rewiring of host cytoskeletal remodeling by SteC. We identified FMNL subfamily formins as the host targets which bound SteC *in vivo* and *in vitro*. We could further show that SteC phosphorylates these formins *in vitro* at S171 (FMNL2; S184 for FMNL1) and at residues in the same functional region, which promote interactions with Cdc42 and thereby actin polymerization (Kühn et al. 2015). Thus, our current model is that SteC directly binds to and phosphorylates FMNL proteins, promoting their interaction with Cdc42 and the recruitment of the complex to the SCV to stimulate actin polymerization (Figure 7F). In agreement with this model, we observed Cdc42 to co-purify with SteC and FMNL1 in macrophages (Figure 2A). However, dominant negative versions of Cdc42 were shown in the past to still allow SPI-2-dependent actin assembly (Unsworth et al. 2004). Further work will be required to elucidate whether SteC has different specificity for different FMNL subfamily members, the molecular events triggered by binding and phosphorylation of FMNLs by SteC, including the level of involvement of Cdc42, and whether the SteC-FMNL binding and regulation are linked to the previously reported modulation of MAPK signaling by SteC (Odendall et al. 2012).

There are many stronger interactions in our study that await further characterization. For example, we found a functional group comprised of the Rab GTPases Rab10, Rab13 and Rab14, targeted by SifB and PipB2 in HeLa cells. Rab10, Rab13 and Rab14 are involved in ER dynamics (English and Voeltz 2013), transport of surface proteins to the cell membrane (Wang et al. 2010; Sano et al. 2008; Sun et al. 2010), tight junction assembly *via* regulation of PKA signaling (Köhler, Louvard, and Zahraoui 2004) and TGN-associated recycling (Nokes et al. 2008; Junutula et al. 2004; Kitt et al. 2008). Furthermore, all three Rab proteins have been linked to various bacterial infectious diseases (Stein, Müller, and Wandinger-Ness 2012), such as intracellular survival of *M. tuberculosis* (Kyei et al. 2006), *and Chlamydia* species (Rzomp et al. 2003). Interrogation of Rab-dependent vesicular trafficking may provide new insights into *S*Tm pathogenesis, especially as several Rab proteins (e.g. Rab29, Rab32) have been implicated in intracellular pathogenicity in the human-adapted pathogen, *Salmonella enterica* serovar Typhi (Spanò, Liu, and Galán 2011; Baldassarre et al. 2019).

In conclusion, we aimed to bridge a technological gap common to host-pathogen PPI studies, which were until now performed exclusively in non-physiological conditions. Our study can be a starting point for more systematic and unbiased studies of host-pathogen PPIs in a native infection context, providing a better understanding of the degree and nature of effector cooperation, which is of high relevance in bacterial pathogens with large effector arsenals. Understanding how pathogens and pathobionts directly modify host pathways *via* secreted proteins will uncover new insights into the diversity and evolution of pathogenicity, as well as provide novel tools and targets to modulate immune responses.

## Acknowledgements

We thank W. Annaert (VIB, KU-Leuven) for the NPC1 knockout cells, B. Bukau (ZMBH Heidelberg) for the GroEL antibody, and F. Kage (Dartmouth Geisel School of Medicine, Hanover, USA) for advice on FMNL2/3 knockout cells. PW was supported by the EMBL International PhD Programme (EIPP) fellowship and JS and CP partially by fellowships from the EMBL Interdisciplinary Postdoc (EIPOD) programme under Marie Sklodowska-Curie Actions COFUND (grant number 291772). KR was supported through intramural funds from the Helmholtz Society. This work was funded by EMBL core funding.

## Author contributions

Conceptualization: AT, JS, PW. Bacterial Strain creation: JS, KF, PW, KS. Proteomics sample preparation: PW, MR, JS, KF. Proteomic data analysis: FS and PW, with input from JS, MMS and AT. Network analysis: PW. Biochemistry and cell Biology: PW, JS, LAK, KS. Kinase assay: CV and PW. Phosphoproteomics: CP. Experimental design: AT, JS, PW, LAK, MG, KR, OSM, DH. Reagent provisions: DH, KR, MG, MMS. Manuscript writing: JS, PW, AT with input from all authors. Figures: PW, JS, LAK, CV, CP. Supervision: AT, JS, DH, MMS & OSM.

## Data availability

All raw AP-QMS and phosphoproteomic data has been deposited at ProteomeXchange (http://www.proteomexchange.org/).

## Code availability

The code and pipelines used for data analysis are available upon request.

## Declaration of interest

The authors declare no competing interests.

## SUPPLEMENTARY FIGURE LEGENDS

**Figure S1.**
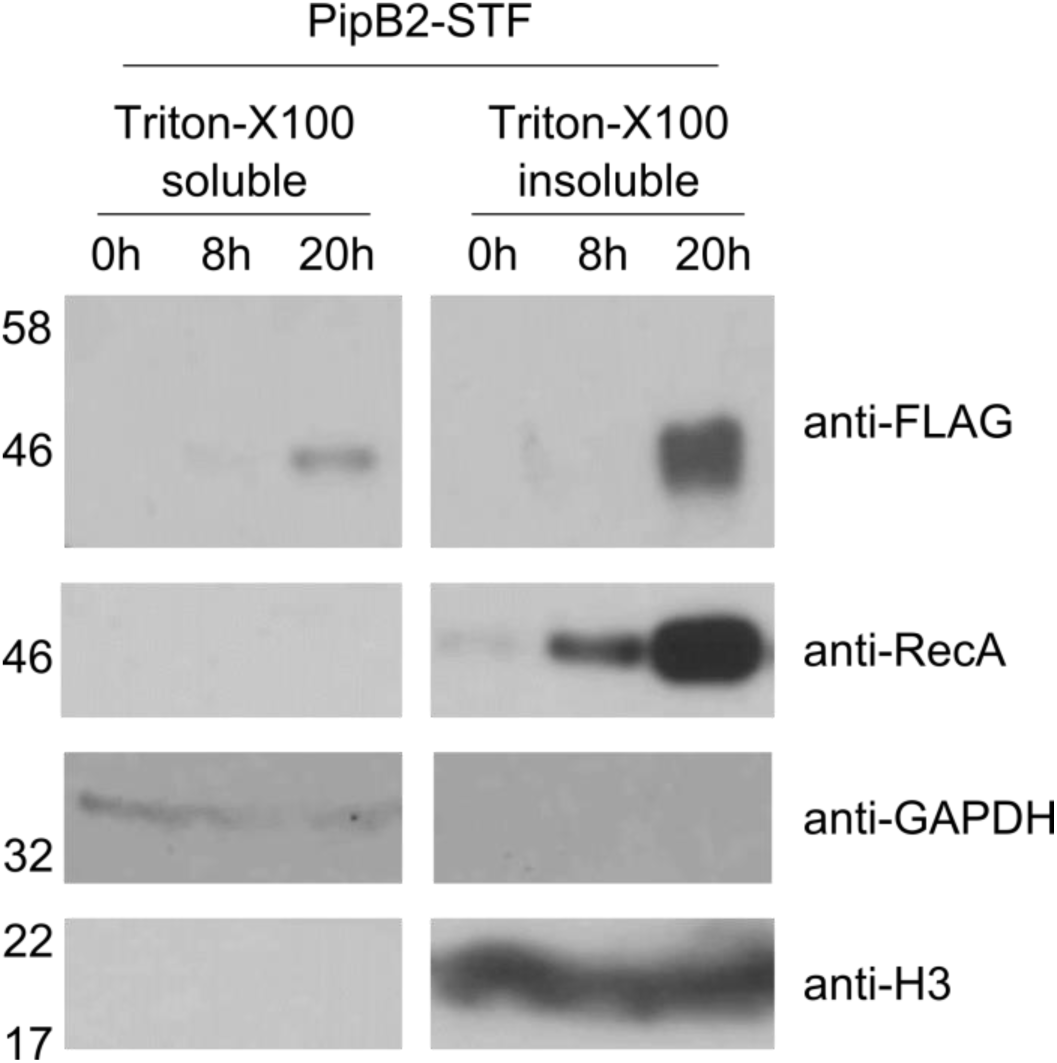
Time-dependent effector expression and translocation of PipB2 probed by Western Blot. RAW264.7 were infected with PipB2-STF expressing *S*Tm. 0 hpi, 8 hpi and 20 hpi, cells were lysed in 0.1 % Triton-X100 and the soluble (cytosolic) and insoluble (nuclei, *S*Tm) fractions were separated by centrifugation. Probing with anti-FLAG antibody shows presence of the effector PipB2-STF in the soluble and insoluble fractions. As loading controls, anti-RecA (bacterial), anti-GAPDH (cytosolic) and anti-H3 (nuclei) were used. Time-dependent expression and translocation was assessed in one experiment.

**Figure S2.**
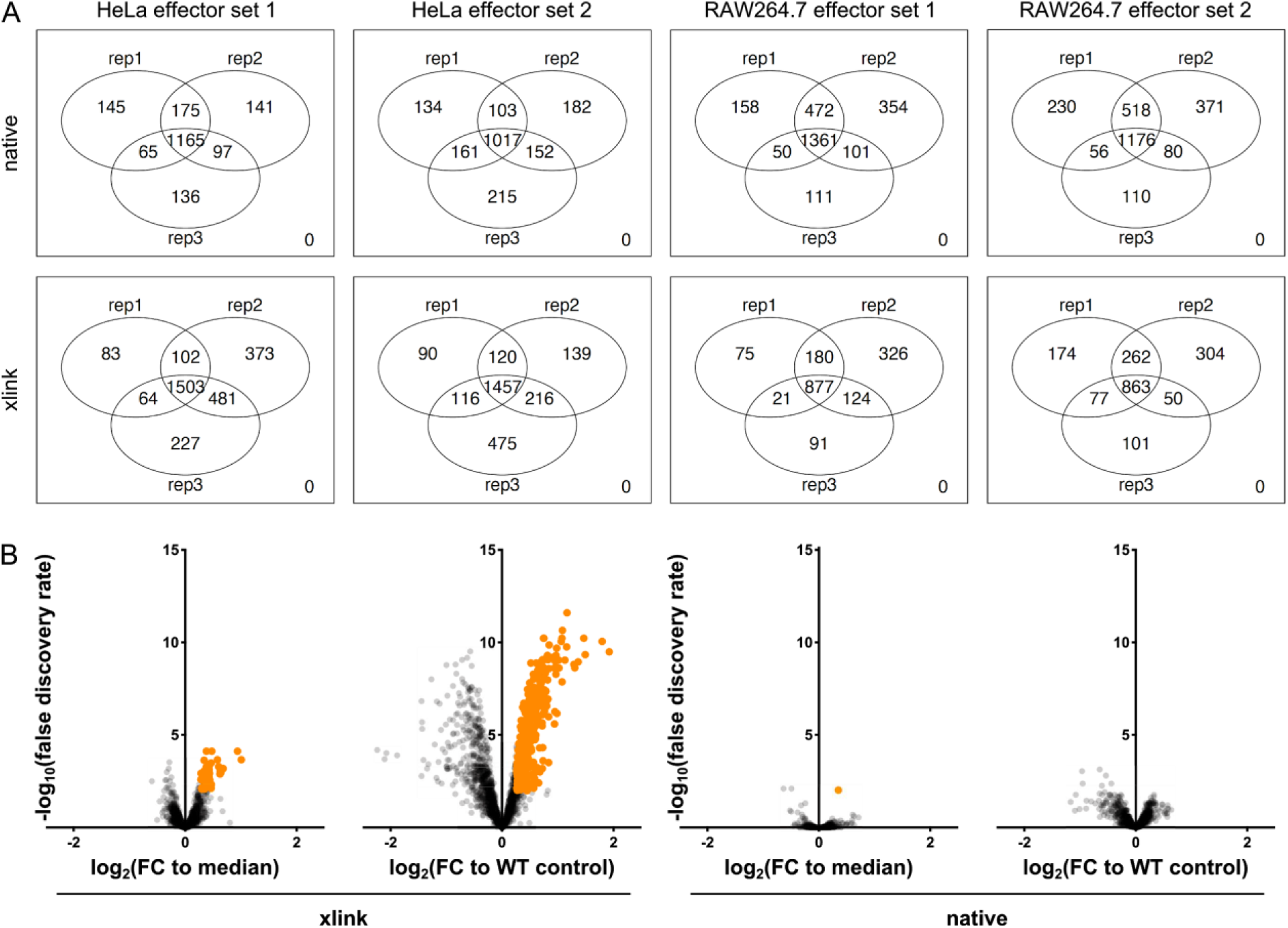
Replicate reproducibility and comparison of normalization to untagged control (WT) with median. (A) Venn diagrams summarizing the number of proteins identified in the different replicates of all TMT10 runs. Effector set 1 and 2 refers to the two 10-plexes (WT + 9 effectors) in which the 20 effectors were split. Only proteins with at least two unique peptides were considered and only those in at least 2 replicates were used for further analysis. (B) Volcano plots showing fold enrichment for targets identified in RAW264.7 after x-linked (left panels) or native (right panels) pulldown of SteE-STF at 20 hpi (as an example). Fold changes (Log_2_) for each protein were calculated by dividing the abundance (signal sum) per TMT channel, per run, by either the median abundance of a given protein across the entire TMT run (first and third panel) or by the abundance of the respective protein in the untagged control (second and fourth panel). Thresholds for hit calling was set to a False Discovery Rate (fdr) of 1% and a Fold Increase of >20% (Fold Change 1.2) (Log_2_). Hits are colored orange and non-hits are displayed in grey. Using normalization with respect to untagged wildtype (WT) control (left 2 panels) displays worse signal-to-noise ratio compared to median normalization (right 2 panels). This is especially true for crosslinked samples.

**Supplementary figure S3.**
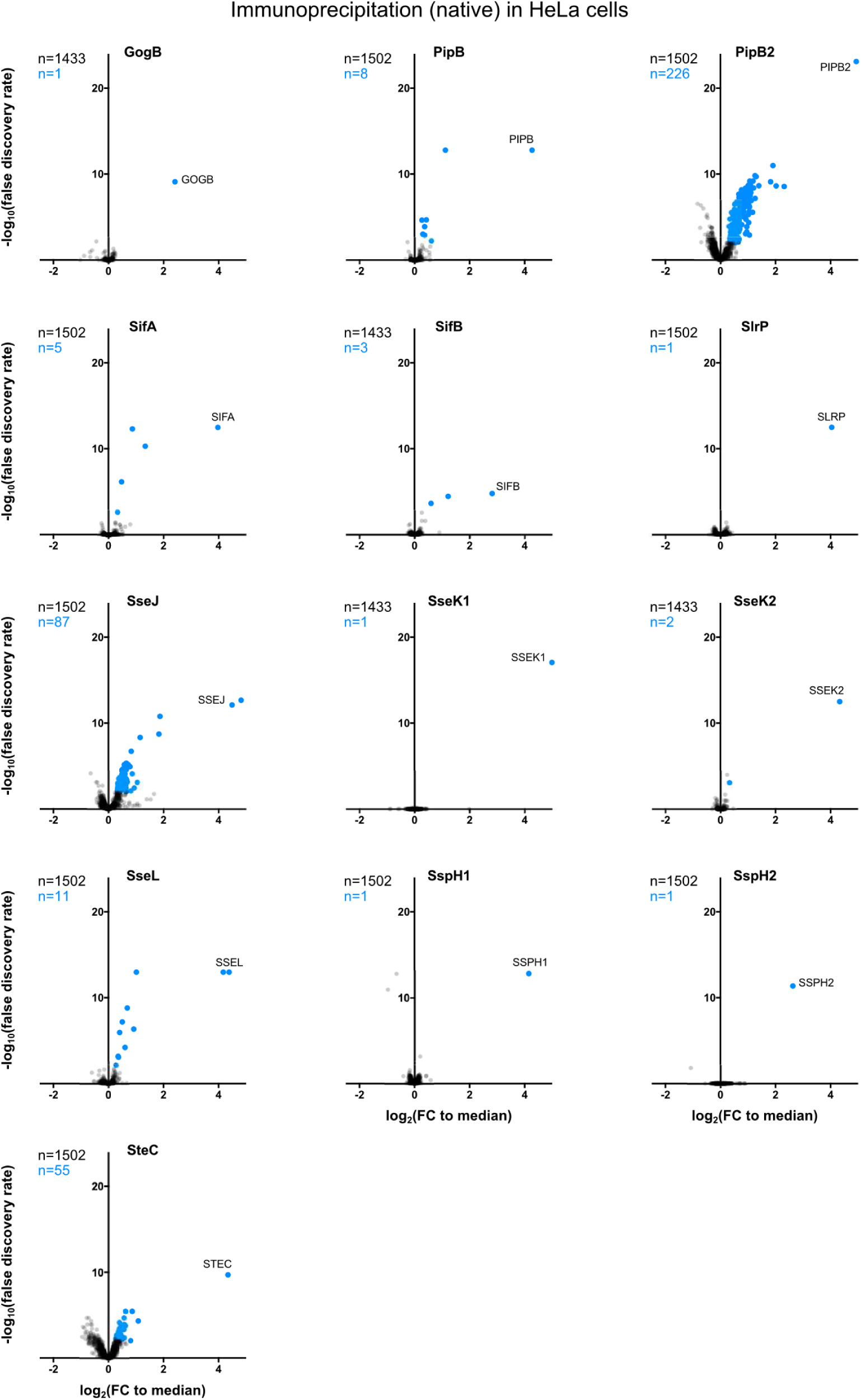
Volcano plots displaying fold enrichment for each protein in HeLa cells after native pulldown at 20 hpi for each tested STF-tagged effector. Fold changes (Log_2_) for each protein were calculated with respect to the median and hits were called as described in Figure S2. Hits are colored blue and non-hits are displayed in grey. See Tables S2 and S3 for all data or hits only for both cell types, respectively.

**Supplementary figure S4.**
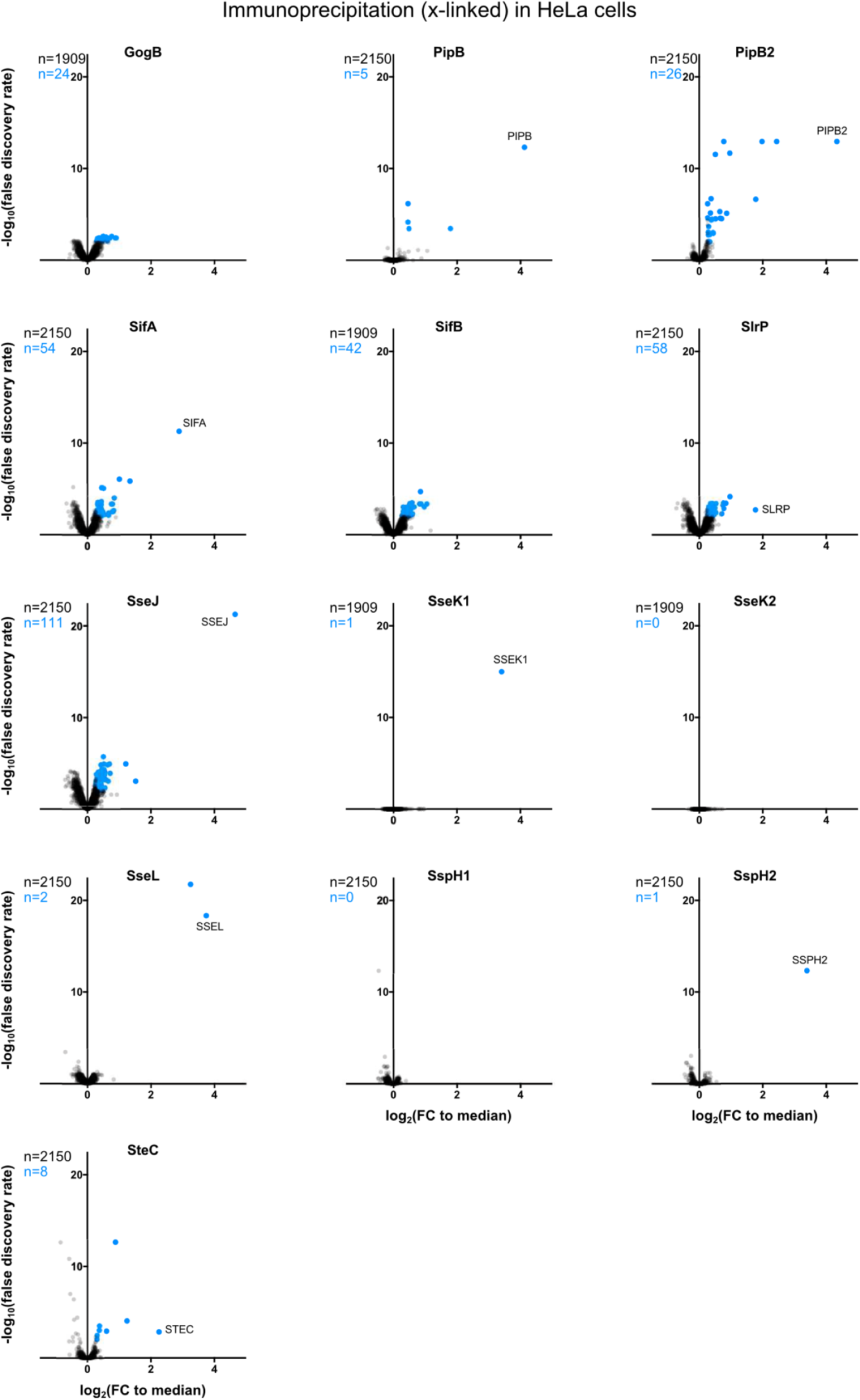
Target enrichment per effector in HeLa cells after cross-linking and pulldown at 20 hpi. Fold changes were calculated and hits called as described in Figure S3, coloring is as in S3. See Tables S2 and S3 for all data or hits only for both cell types, respectively.

**Supplementary figure S5.**
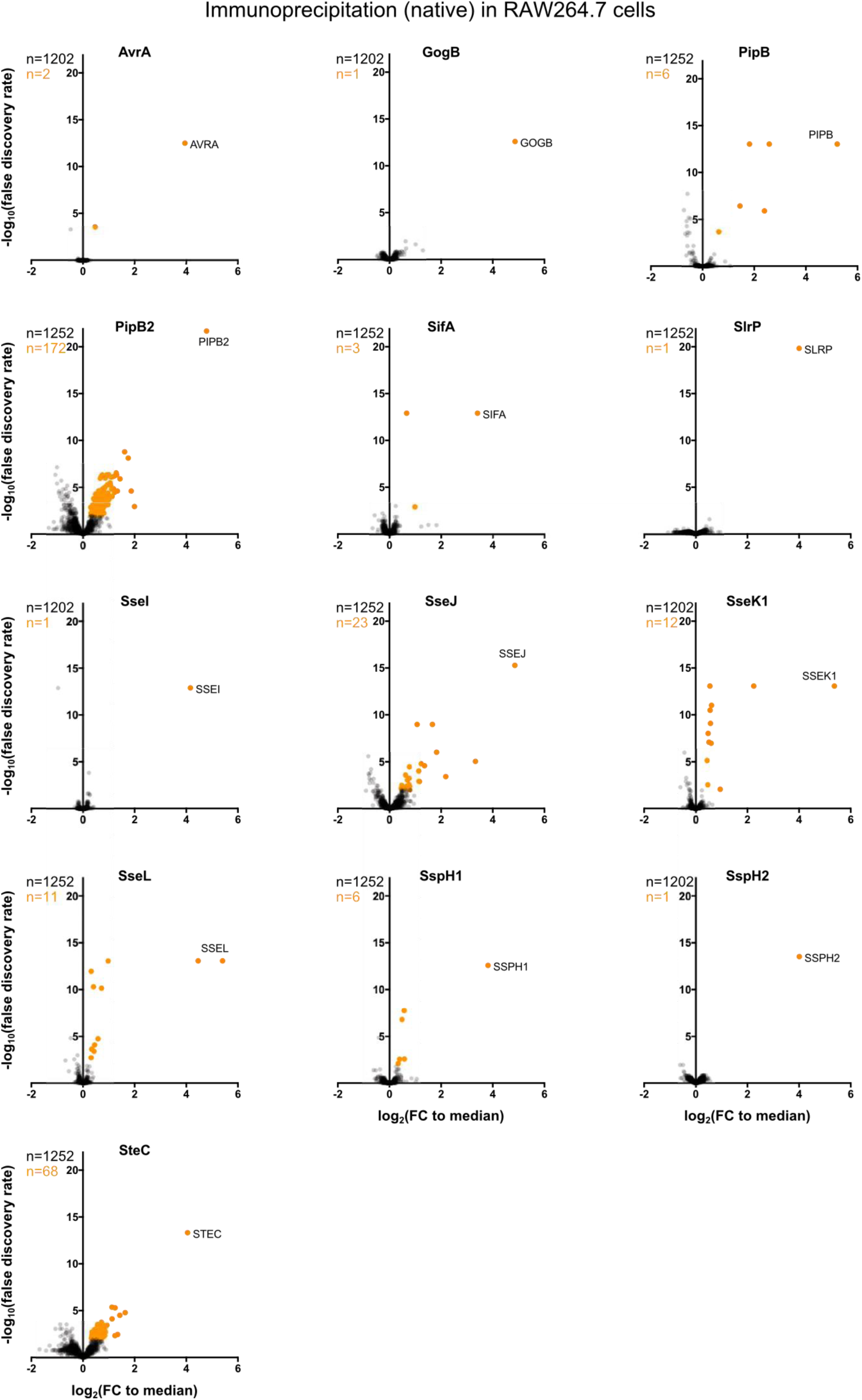
Target enrichment per effector in RAW264.7 cells after native pulldown at 20 hpi. Fold changes were calculated and hits called as described in Figures S3 and S4. Hits are colored orange and non-hits are displayed in grey. See Tables S2 and S3 for all data or hits only for both cell types, respectively.

**Supplementary figure S6.**
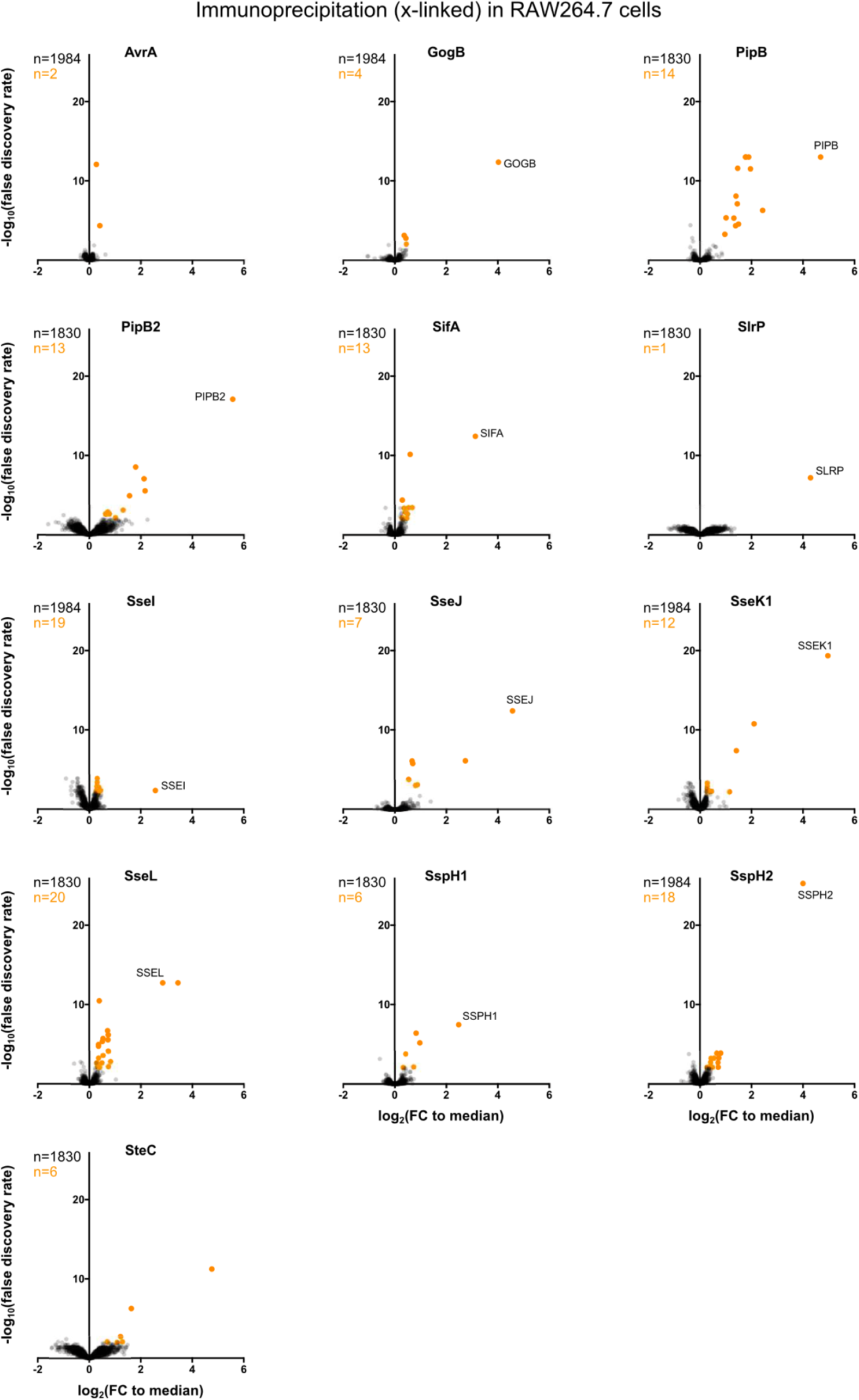
Target enrichment per effector in RAW264.7 cells after cross-linking and pulldown at 20 hpi. Fold changes were calculated and hits called as described in Figures S3 - S5, hits are colored as in S5. See Tables S2 and S3 for all data or hits only for both cell types, respectively.

**Figure S7.**
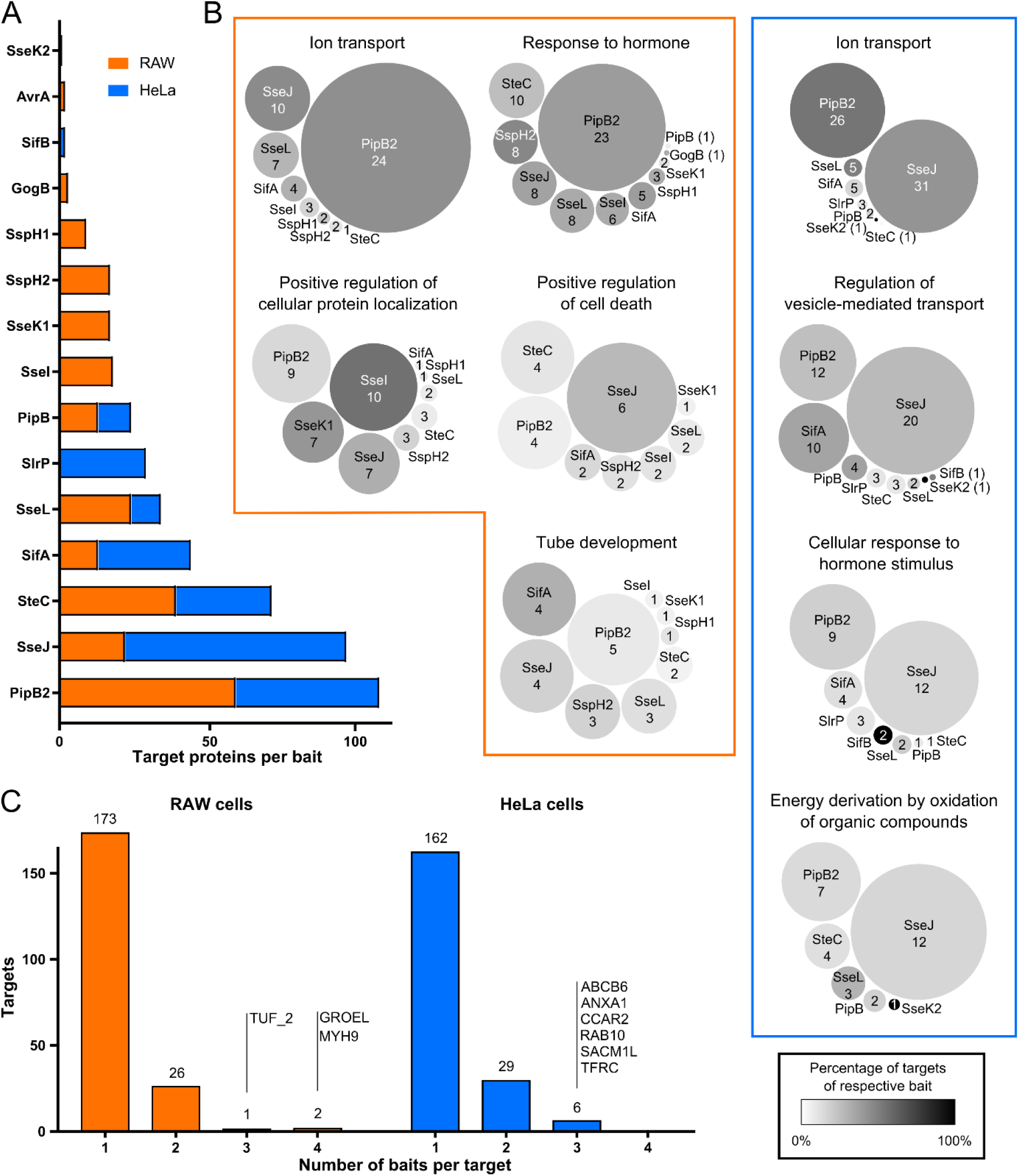
Connectivity within PPI networks. (A) Number of target proteins interacting with each effector in RAW264.7 (orange) and HeLa (blue) cells, ordered by total number of protein targets. (B) Effectors affecting various GO terms. Bubble size corresponds to the percentage of targets associated with a given GO-term interacting with the respective effector (number of eukaryotic protein targets is indicated). Shade of the bubble corresponds to the percentage of target proteins associated with any given GO-term with respect to the total number of proteins interacting with the respective effector (color as indicated in the spectrum). (C) Histogram of the number of *S*Tm effector proteins (baits) interacting with each target protein in RAW264.7 (left side, orange) and HeLa cells (right side, blue). Most targets interact with a single bait, but several can work as connection points between different effectors. Names are indicated for targets with more than 3 PPIs with effectors.

**Figure S8.**
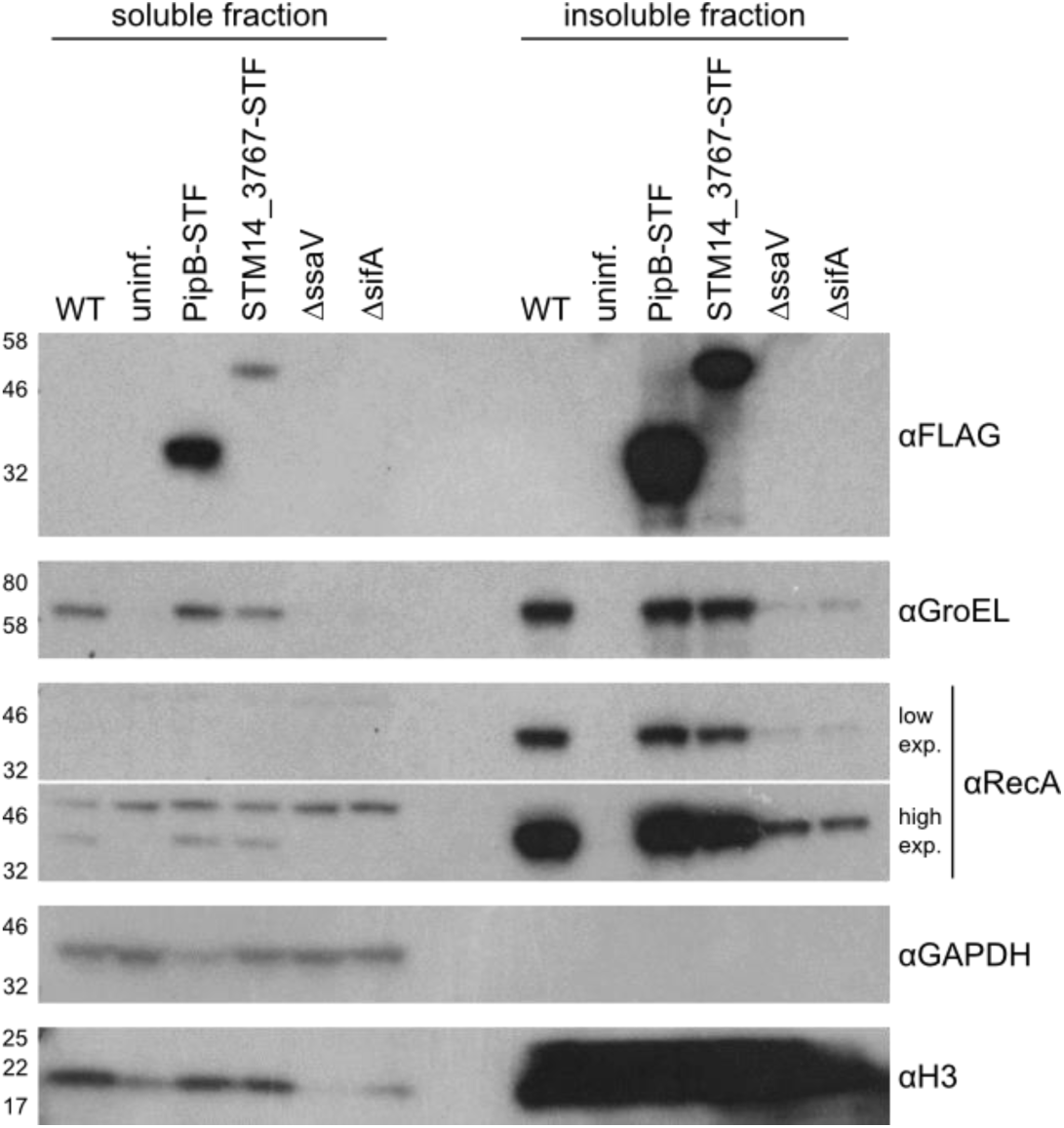
Subcellular fractionation showing GroEL enrichment in the host cell cytoplasm. RAW264.7 macrophages were infected with wildtype *S*Tm, several tagged stains, a Δ*sifA* mutant (which displays decreased vacuolar stability and less proliferation in macrophages) and a Δ*ssaV* mutant (T3SS2 deficient). Western blot was performed after harvesting at 20 hpi in Triton-X100 containing lysis buffer in a single replicate. The soluble fraction (cytoplasm) is displayed on the left side, the insoluble fraction (*S*Tm, nuclei) on the right. αFLAG antibody was used to determine translocation of tagged effectors and αGroEL to determine presence of GroEL in the respective fractions, loading controls: αRecA (bacterial), αGAPDH (cytoplasmic fraction), αH3 (nuclear). In addition to the presence of the effector protein, PipB, in the soluble fraction (previously described), we also saw the bacterial proteins GroEL and STM14_3767, a bacterial itaconate CoA-transferase which interacted with PipB in the host cytoplasm, yet not the bacterial loading control RecA.

**Figure S9.**
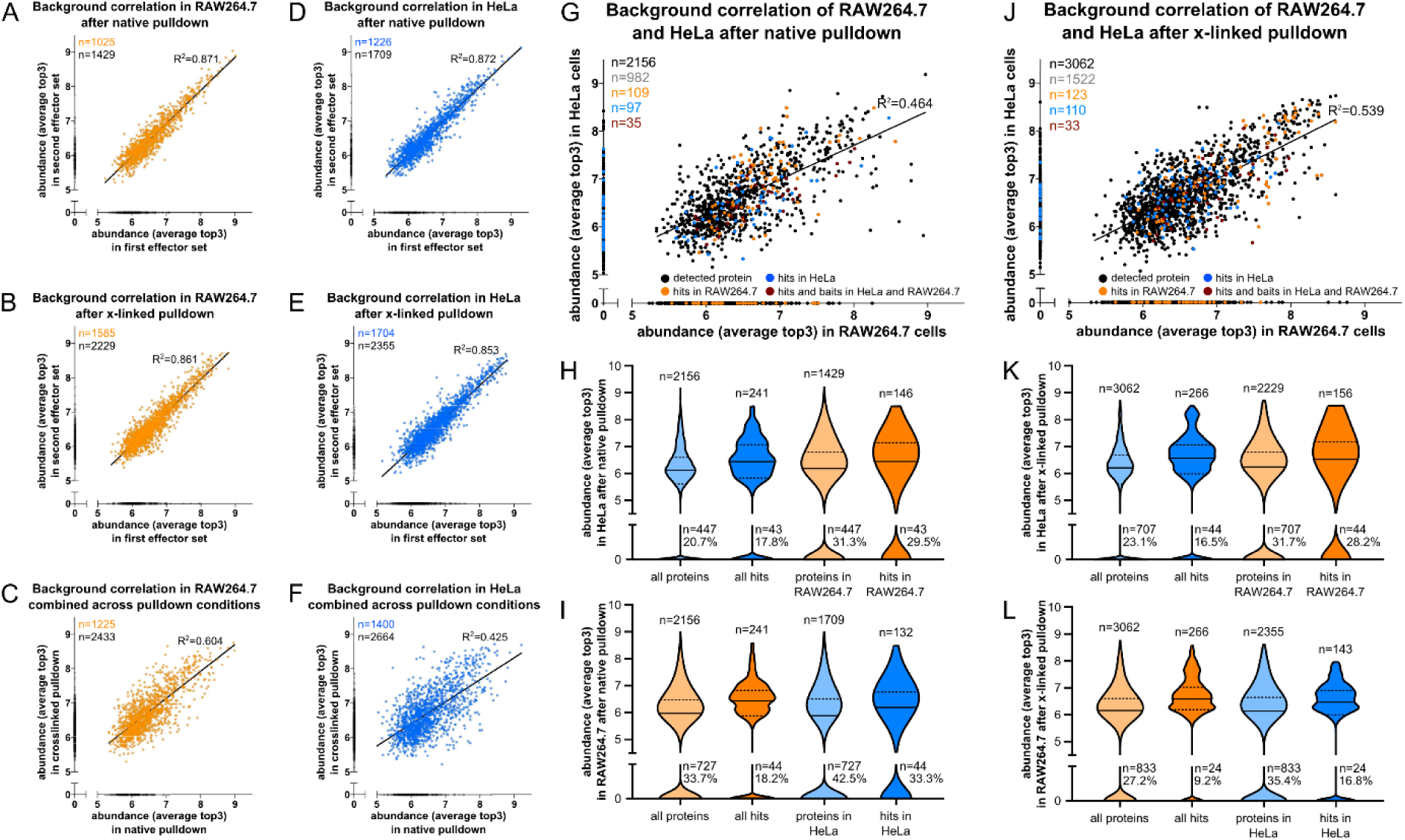
Comparison of RAW264.7 and HeLa expression protein expression. Orthologs were found based on protein name and using the OMA-browser (Altenhoff et al. 2018), and if no ortholog was found, or if the protein was not detected in the other cell line, abundance was set to 0 – more detailed information is provided in Experimental Procedures. Correlations of protein abundances in various runs, as indicated in the title of each respective scatter plot. (A-C) Orange (RAW264.7) and (D-F) blue (HeLa) blots from top to bottom: (A, D) batch comparison in native pulldown; (B, E) batch comparison in pulldown after cross-linking; (C, F) average of native pulldowns vs. average of crosslinked pulldowns. Large scatter plots: (G) Native pulldown in RAW264.7 vs. HeLa. Hits are indicated: hits in both cell lines in red, hits in HeLa cells in blue, hits in RAW264.7 in orange. (H, I) Violin plots as in Figure 4C summarizing the protein abundance in HeLa cells (top) and RAW264.7 macrophages (bottom) after native pulldown, i.e. quantification of the x- and y-axis, as well as blue and orange dots of the summarizing scatterplot. (J) cross-linked pulldown in RAW264.7 *vs* HeLa, hits are annotated as in panel G. (K, L) Violin plot summarizing panel J, quantification and display as in panels H and I, respectively.

**Figure S10.**
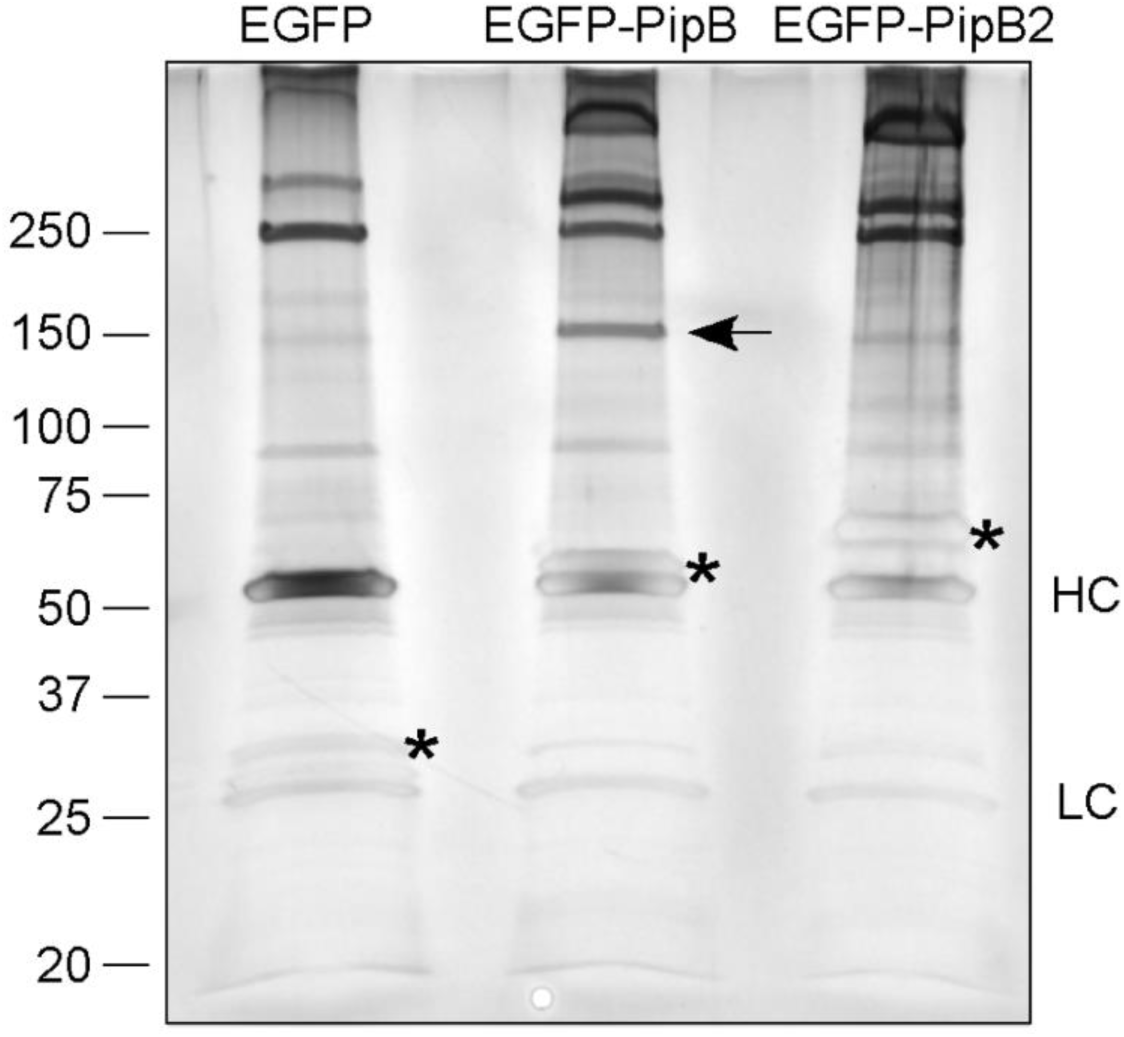
Interaction of PipB, but not PipB2, with PDZD8 after transfection. SilverQuest Silver stain of proteins that have been co-IPed from HeLa cells with either EGFP, EGFP-PipB or EGFP-PipB2. Protein indicated by arrow in EGFP-PipB lane was sent for LC-MS/MS analysis and identified as PDZD8. Asterisks denote EGFP and EGFP-fusion proteins that were immunoprecipitated in each condition. HC, heavy chain; LC, light chain.

**Figure S11.**
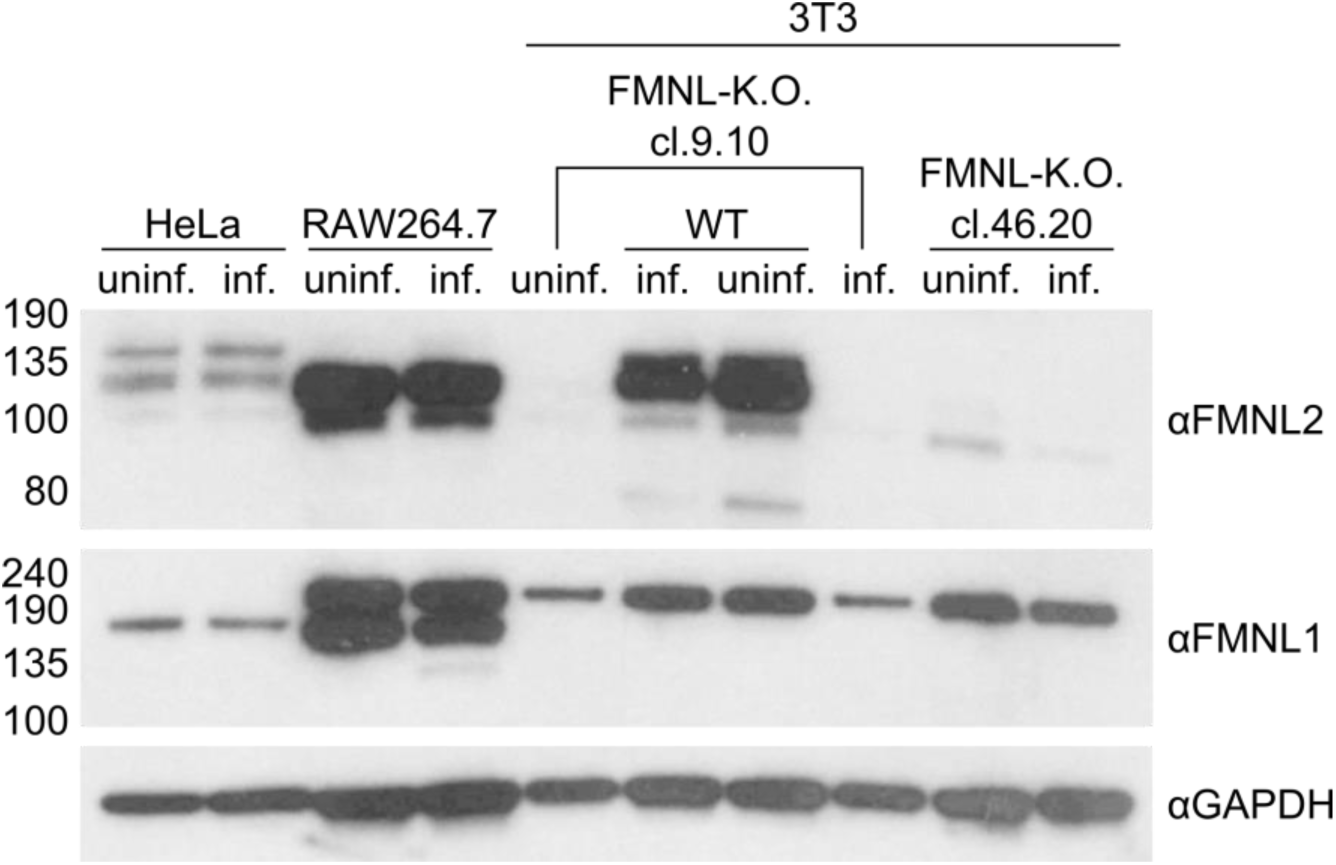
Expression of FMNL1, FMNL2 and FMNL3 in all cell types used for this study. Western Blot (single replicate) showing the presence of FMNLs in the various cell lines used. Protein detection with αFMNL1, αFMNL2 (which also cross-reacts with FMNL3, (Kage, Winterhoff, et al. 2017) and αGAPDH antibodies was performed as described in the Experimental Procedures section. Cleared cell lysate (Triton-X100 soluble fraction) was loaded in all cases. Infected samples (inf.) are at 8 hpi.

### Supplemental information file list

**Figure S1**. STm effector expression and translocation timeline

**Figure S2**. Replicate reproducibility and fold-change calculation

**Figure S3-6**. Volcano plots of all TMT10-runs (S3: HeLa native, S4: HeLa: crosslinked, S5: RAW264.7 native, S6: RAW264.7 crosslinked)

**Figure S7**. Connectivity within PPI networks

**Figure S8**. Subcellular fractionation showing GroEL enrichment in the host cell cytoplasm

**Figure S9**. Correlation of RAW264.7 *vs* HeLa expression data

**Figure S10**. Interaction of PipB, but not PipB2 with PDZD8 after transfection

**Figure S11**. Expression of FMNL1, FMNL2 and FMNL3 in all cell types used for this study

**Table S1**. STm tagged effector library; list of tagged effectors and performance

**Table S2**. STm-host Protein-protein interactions in RAW264.7 and HeLa cells. Summary of all Limma results from AP-QMS work. (One sheet cell line and pulldown condition)

**Table S3**. STm-host Protein-protein interactions in RAW264.7 and HeLa cells. Hits from AP-QMS. (One sheet for RAW264.7 macrophages and one for HeLa cells)

**Table S4**. Summary of previously published PPIs during STm infection that were identified in our AP-QMS approach

**Table S5**. Functional interactions extracted from STRING DB version 11 to build the interaction networks

**Table S6**. List of all GO-terms and GO clusters identified in RAW264.7 and HeLa cells.

**Table S7**. Protein abundances in RAW264.7 and HeLa cells. Entire dataset. (Protein abundances as detected in all different conditions, combined into one with pulldown condition and cell line in the various columns)

**Table S8**. List of PPIs chosen for validation with reciprocal pulldown, including information on log(FC) in the AP-QMS work, feasibility of target pulldown and detectability of reciprocal interaction

**Table S9**. Limma results of separate TMT10-run focusing on SseJ (crosslinked pulldown 20 hpi in HeLa cells)

**Table S10**. Identified phosphosites in *in vitro* kinase assay of SteC with N-terminal regions of FMNL1 and FMNL2

### Key Resources Tables

**Table S11**. List of primers used

**Table S12**. List of bacterial strains used

**Table S13**. List of plasmids used

**Table S14**. List of antibodies used

## Experimental procedures

No statistical methods were used to predetermine sample size.

### Media, chemicals and reagents

The following chemicals and reagents used were purchased from Sigma: DMSO (cat. nr. D8418), Triton-X100 (x100), heat inactivated Fetal Bovine Serum (FBS) (F9665-500ML), Phalloidin ATTO-647N (65906), gentamicin (G1914); Gibco: DMEM 4.5 g/L glucose (41965); Roche cOmplete mini EDTA-free protease inhibitors (11873580001); Life Technologies Hoechst 33342 (H3570); Thermo Scientific Pierce™ formaldehyde 16% (w/v) (28908). Antibodies are listed in table S14 including the distributor and catalog number. Bacterial antibiotic selection was performed on LB agar containing ampicillin 100µg/mL or 30µg/mL kanamycin at 37**°**C.

### Bacterial strains and plasmids

All strains used in this study are listed in Table S12. *Salmonella enterica* subsp. Typhimurium 14028s (*S*Tm) wildtype was used to generate the tagged effector library as described below. Single gene deletion mutants (Δ*sseJ*, Δ*sseL*, Δ*steC*, Δ*prgK*, Δ*ssaV*, Δ*sifA*) were struck out from a single deletion collection (Porwollik et al. 2014) followed by PCR confirmation and retransduction into the wildtype *S*Tm 14028s background using P22 phage. To generate the Δ*sseJ*Δ*sseL* double mutant, FLP-FRT mediated excision of the antibiotic resistance cassette was performed as previously described (Datsenko and Wanner 2000), followed by P22 transduction of the second mutated loci. Resulting double mutants were verified by colony PCR. *S*Tm SL1344 wildtype, Δ*pipB* and Δ*pipB2* strains have been described previously (Hoiseth and Stocker 1981; Knodler et al. 2002, 2004). The complementing plasmids, pPipB-2HA and pPipB2-2HA, are pACYC184 derivatives and have been described previously (Knodler et al. 2002, 2004).

The *S*Tm 14028s tagged effector library was generated as follows. To generate the template plasmid (pJPS1) we cloned the 2xSTREP-TEV-3xFLAG (STF) tag into the MCS (EcoRI-HindIII) of pQE30 and designated pMZ2. The pMZ2 plasmid was then used as a PCR template to amplify a 2xSTREP-TEV-3xFLAG tag together with the pKD4 kanamycin resistance cassette using primers JPS26 and JPS27. This amplicon was then T/A cloned into pGEM®-T Easy according to manufacturer’s instructions followed by sequence verification and designation as pJPS1. Purified pJPS1 was used as template DNA to amplify and introduce a 2xSTREP-TEV-3xFLAG (STF) tag followed by a kanamycin resistance cassette at the C-terminus of chromosomally encoded genes via λ-red recombinase (Datsenko and Wanner 2000; Uzzau et al. 2001). Clones were selected on LB agar containing kanamycin 30µg/mL and verified PCR and sequencing of the C-terminal region of the targeted gene. The resulting tagged effectors expressed the following C-terminal STF affinity tag sequence; GGAAAG<u>WSHPQFEK</u>GGGSGGGSGGGS<u>WSHPQFEK</u>GENLYFQGA<u>DYKD</u>HDG<u>DYKD</u>HDI <u>DYKDDDDK.</u> See Table S1 and S12 for the complete list of effectors targeted.

To avoid disturbing the C-terminal prenylation motif of the effector *sifA*, the STF tag was chromosomally inserted within the open reading frame between residues D136 and I137 using a two-step selection method related to λ-red recombination was (Kolmsee and Hengge 2011). Briefly, to generate an *S*Tm 14028s strain amenable to pKD45 two-step selection, the endogenous *S*Tm *ccdAB* locus (STM14_5550 and STM14_5550) was deleted via λ-red recombination (Datsenko and Wanner 2000; Uzzau et al. 2001) using primers JPS38 and JPS39, followed by PCR verification and P22 transduction to a wildtype background and designated *S*Tm Δ*ccdAB::Cm*. A fragment of the plasmid pKD45 (Datsenko and Wanner 2000) encoding a kanamycin-resistance cassette and a *ccdB* toxin under the control of a rhamnose-inducible promoter was amplified using primer pairs JPS14 and JPS15 containing extensions homologous to the *sifA* locus (STM14_1400). The resulting amplicon was chromosomally integrated into *S*Tm Δ*ccdAB::Cm* using λ-red recombination and selected on LB agar containing 30µg/mL kanamycin (Datsenko and Wanner 2000). Positive *sifA*::*Kan*-*ccdB* transformants were verified by PCR and tested for L-rhamnose sensitivity on M9 minimal agar. Using primers JPS28 and JPS29 and the pJPS1 plasmid as DNA template, an amplicon containing overhangs with sequence homology to *sifA* and an internal STF sequence was amplified and integrated onto the chromosome using λ-red recombinase (Datsenko and Wanner 2000). Transformants were counter-selected on M9 minimal agar containing 0.5% L-rhamnose after incubation at 30°C for 2 days and verified by PCR. A list of STF-tagged effectors generated is listed in Table S1, along with summarized test expression behavior in both HeLa and RAW264.7 cells.

For ectopic expression in mammalian cells, the *pipB* open reading frame was amplified from *S*. Typhimurium SL1344 genomic DNA with the oligonucleotides pipBGFP-N5’ and pipBGFP-N3’2. The amplicon was digested with BglII/SalI and ligated into BglII/SalI-digested pEGFP-C1 (Clontech) to create EGFP-PipB. EGFP-PipB(Δ270-291) was created by amplification with pipBGFP-N5’ and GFPPipB-269R, digestion with BglII/SalI and ligation into pEGFP-C1. EGFP-PipB2 has been described previously (Knodler and Steele-Mortimer 2005). PDZD8 was tagged at the C-terminus with a myc epitope for immunodetection. The coding sequence plus an upstream Kozak sequence were amplified from a PDZD8 cDNA clone, MGC:27107 IMAGE:4837939 (The CCSB Human ORFeome Collection) with the oligonucleotides PDZK8-EcoRI-Kozak and NM_173791-Xho. The amplicon was ligated in EcoRI/XhoI-digested pCMV-Tag 5A (Stratagene) to create pKozak-PDZD8-myc.

For expression in yeast, EcoRI/BglII fragments encoding full length, residues 1-281, residues 1-271, residues 1-188 and residues 189-291 of PipB were amplified from SL1344 genomic DNA with the following oligonucleotide pairs and ligated into pGAD424 (Clontech): pGAD-PipB-1F and pGAD-PipB-291R, pGAD-PipB-1F and pGAD-PipB-281R, pGAD-PipB-1F and pGAD-PipB-271R, pGAD-PipB-1F and pGAD-PipB-188R, pGAD-PipB-189F and pGAD-PipB-291R. Full-length and fragments of PDZD8 were PCR amplified as EcoRI/SalI fragments from PDZD8 cDNA (details above). Amplicons were digested and ligated into pGBT9 (Clontech). The following oligonucleotide pairs were used: pGBT9-PDZK8-F and pGBT9-PDZK8-R for pGBT9-PDZD8, pGBT9-PDZK8-F2 and pGBT9-PDZK8-R for pGBT9-PDZD8(Δ1-338), pGBT9-PDZK8-F and pGBT9-PDZK8-R2 for pGBT9-PDZD8(Δ930-1154). Overlap extension PCR (Horton et al. 1989) was used to create pGBT9 constructs that were deleted for residues 368-461 (pGBT9-PDZD8ΔPDZ), residues 494-814 (pGBT9-PDZD8(Δ494-814)) and residues 841-887 (pGBT9-PDZD8ΔC1).

### Cell culture conditions

RAW264.7 macrophages (ATCC TIB-71) and HeLa epithelial cells (ATCC CCL-2) were cultured at 37°C, 5% CO_2_ in DMEM containing 4.5g/l glucose (Gibco). Cells were passaged at 90% confluency and were not used beyond passage number 15. For cell passaging and seeding, media was removed, cells were washed once in pre-warmed PBS and detached by incubation in 0.05% trypsin-EDTA (for HeLa cells, Thermo Fisher, cat. Nr. 25300054) or accutase (for RAW264.7 cells, Thermo Fisher, cat. Nr. A1110501) at 37°C for ∼3 min. Complete media was added to the cell suspension and cells were counted using trypan blue staining in a Biorad TC20 automated cell counter. If cells were prepared for infection, the following cell numbers were seeded 20h prior to the infection: For 96-wells (Zell-Kontakt, cat. Nr. 21315241), 7.5×10^3^ HeLa and 3×10^4^ RAW264.7 cells; 6-wells (Thermo Scientific, cat. Nr. 10119831), 2×10^5^ HeLa and 9×10^5^ RAW264.7 cells; 15cm dishes (Greiner, cat. Nr. 639160), 3.5×10^6^ HeLa and 15.4×10^6^ RAW264.7 cells. For large-scale AP-QMS experiments, five 15cm dishes were seeded per effector per condition, equaling a total cell number of ∼75×10^6^ and 17.5×10^6^ for cells for RAW264.7 and HeLa cells, respectively. NIH 3T3 wildtype and derived FMNL2/3 double knockout clones 9.10 and 46.20 were maintained as described before (Kage, Steffen, et al. 2017). HeLa cells harboring an NPC1 knockout were maintained as previously described (Tharkeshwar et al. 2017).

### Infection of RAW264.7 macrophages and HeLa cells

For infection of RAW264.7, *S*Tm strains were cultured overnight at 37°C, washed in PBS and added to the cells at a multiplicity of infection (MOI) of 100. For infections carried out in multiwell plates, the bacteria were spun down at 170G for 5 min to increase contact between bacteria and macrophages. The infection was performed for 30 min at 37°C, after which the media containing bacteria was removed by aspiration, cells were washed once in pre-warmed PBS. Subsequently, cells were cultured at 37°C in DMEM (4.5 g/l glucose) containing 100µg/ml gentamycin to kill all remaining extracellular bacteria. After 1 hr, the media was replaced with DMEM containing 16 µg/ml gentamycin for the remainder of the experiment (this also denotes time point zero). For HeLa cell infection, overnight cultures of *S*Tm strains were subcultured (300µL overnight culture in 10 ml LB Lennox containing adequate antibiotics) and cultured for 3.5 hr at 37°C in 100 ml Erlenmeyers at 45 rpm (adapted from (Steele-Mortimer 2008)). For infection, a MOI of 100 was used and the infection and gentamicin protection assay were performed as described for macrophages in the previous paragraph. DMEM (1g/l Glucose) was used as growth medium.

### Proteomic sample preparation for AP-QMS

For native harvesting, cells were washed twice in PBS at RT and lysis buffer (PBS, containing 0.1% Triton-X100 and 1x Protease Inhibitor (cOmplete EDTA free, Roche) was added (300µL for 6-well plates, 5ml for 15cm dish). Cells were put at 4°C for 30 min while shaking gently and subsequently scraped off and resuspended by pipetting. The cell lysate suspension was centrifuged at 4°C for 15 min at 20,000G to clear the lysate. A small sample of the cleared lysate was saved as “Total” sample, the remaining lysate was directly used for immunoprecipitation. For harvesting after crosslinking, the cells were washed twice in PBS at RT and crosslinking buffer (PBS, containing 1mM DSP (Thermo Fisher, cat. nr. 22585)) was added. Crosslinking was performed for 2 hr at 4°C and quenched using 20mM Tris-Cl at pH Cells were washed twice in quenching buffer and subsequently subjected to the lysis protocol described above.

For pulldown of tagged *S*Tm effectors, anti-FLAG M2 affinity gel (Sigma, A2220) was used. 50µL of the slurry per sample were washed twice in lysis buffer (centrifugation for 1 min at 4°C and 5000rpm). The beads were added to fresh, cleared lysate and incubated for at least 4h (native samples) or O.N. (x-linked samples) at 4°C while tumbling. After bead incubation, the suspension was spun down at 4000rpm for 10 min (4°C) and the supernatant was removed. The beads were washed four times in 1ml cooled washing buffer (PBS containing 0.01% Triton-X100), using centrifugation at 5000rpm for 1 min (4°C) for sedimentation. After the final wash, all remaining buffer was removed, and 40µL elution buffer (PBS containing 150µg/ml 3x FLAG peptide and 0.05% RapiGest) was added. After 1h overhead tumbling at 4°C, the suspension was spun down at 8200rpm at 4°C and the supernatant was removed. 40µL elution buffer were added and one more round of elution was performed.

### TMT-labeling of AP-QMS samples and Mass Spectrometry

Within each TMT-10plex, untagged control (wildtype), as well as 9 STF-effector strains were assessed in parallel (RAW264.7 run 1: WT, PipB, PipB2, SifA, SseJ, SseL, SspH1, SteC, SlrP, run 2: WT, AvrA, GogB, SipB, SpvC, SseI, SseK1, SspH2, SteA, SteE; HeLa run 1: WT, PipB, PipB2, SifA, SseJ, SseL, SspH1, SspH2, SteC, SlrP, run 2: WT, AvrA, GogB, SifB, SpvC, SseF, SseI, SseK1, SseK2, SteA). For each run, all STF-tagged effector strains were seeded and infected at the same time. Prior to MS, 1µL of the elution fractions were used in Western Blot to validate the presence of the effector bait. Total protein concentration was determined using the Pierce Micro BCA kit, according to the manufacturer’s protocol. All samples were adjusted to 10 µg protein in 50 µL volume and were subsequently submitted to the EMBL Proteomics Core Facility. After reduction of disulfide bridges using 10 mM dithiothreitol at 56°C for 30 min in HEPES buffer (50 mM HEPES, pH 8.5), alkylation was performed using 20 mM 2-chloroacetamide at room temperature in HEPES buffer for 30 min under exclusion of light. Samples were prepared according to the SP3 protocol (Hughes et al. 2019) and trypsinized (sequencing grade, Promega, enzyme to protein ratio 1:50) overnight at 37°C. Subsequently, peptides were recovered in HEPES buffer by collecting supernatant on magnet and combining it with a second elution wash of the magnetic beads with HEPES buffer. Peptides were labelled with TMT10plex (Werner et al. 2014) Isobaric Label Reagent (ThermoFisher) according the manufacturer’s instructions. In short, 0.8mg reagent was dissolved in 42 µL acetonitrile (100%) and 4 µL of this stock were added to the samples and incubated for 1 h at room temperature. The reaction was then quenched with 5% hydroxylamine for 15 min. Samples were pooled for the TMT-10plex and then further cleaned using OASIS® HLB µElution Plate (Waters). Subsequently, offline high pH reverse phase fractionation was performed on an Agilent 1200 Infinity high-performance liquid chromatography system, using a Gemini C18 column (3 μm, 110 Å, 100 x 1.0 mm, Phenomenex) with 20 mM ammonium formate (pH 10.0) and 100% acetonitrile as mobile phase (Reichel et al. 2016). The first and two last fractions were discarded prior to LC-MS analysis.

### AP-QMS Data acquisition

Samples were analyzed on an UltiMate 3000 RSLC nano LC system (Dionex) using a µ-Precolumn C18 PepMap 100 trapping cartridge (5µm, 300 µm i.d. x 5 mm, 100 Å) and a nanoEase™ M/Z HSS T3 column 75 µm x 250 mm C18 as analytical column (1.8 µm, 100 Å, Waters). After trapping with a constant flow of 0.1% formic acid in water at 30 µL/min onto the trapping column for 6 min, elution was carried out via the analytical column at a constant flow of 0.3 µL/min with increasing percentage of solvent (0.1% formic acid in acetonitrile): from 2% to 4% in 4 min, from 4% to 8% in 2 min, then 8% to 28% for a further 96 min, and finally from 28% to 40% in another 10 min. The analytical column was coupled to QExactive plus (Thermo) mass spectrometer and mass-spec was performed according to previously described parameters (Perez-Perri et al. 2018).

### AP-QMS Data analysis

IsobarQuant (Franken et al. 2015) and Mascot (v2.2.07) were used to process the acquired data. Peptide search was performed against a Uniprot *Homo sapiens* proteome database (UP000005640, for HeLa cell samples) or a Uniprot *Mus musculus* database (UP000000589, for RAW264.7 cell samples), combined with the *Salmonella* typhimurium (strain 14028s / SGSC 2262) (UP000002695) database containing common contaminants and reversed sequences. The following modifications were included in the search parameters: Carbamidomethyl (C) and TMT10 (K) as fixed modifications, acetyl (protein N-terminus), oxidation (M) and TMT10 (N-terminal) as variable modifications. Mass error tolerance was set as follows: 10ppm for the full scan (MS1) and 0.02Da for MS/MS (MS2) spectra. In addition, a maximum of two missed cleavages were allowed for trypsin, a minimum peptide length of seven amino acids was required and the false discovery rate (fdr) on peptide and protein level was set to 0.01. The output files of IsobarQuant (Franken et al. 2015) were analyzed using the R programming language (ISBN 3-900051-07-0). Only proteins that were quantified with at least two unique peptides and identified in at least two out of three biological replicates were kept for further analysis. The ‘signal_sum’ columns were first annotated to their biological conditions and then a median across all conditions per replicate was computed for each protein. First, potential batch-effects were removed using the ‘removeBatchEffect’ function of the limma package (Ritchie et al. 2015). Second, data were normalized with a variance stabilization normalization (vsn – (Huber et al. 2002)). Finally, missing values were imputed using the impute function (method = “knn”) of the Msnbase package (Gatto and Lilley 2012). Limma was employed again to test for differential expression. Fold changes with respect to the median of the respective run were calculated for each protein in each pulldown. T-values from the limma output were pasted also into fdrtool (Strimmer 2008) in order to compute alternative fdrs. In case the standard deviation of the t-values deviated from 1 to a degree that no convergence of statistically significant hits was observed, the q-values from the fdrtool output were used as alternative fdrs. A protein was annotated as a ‘hit’ with an fdr smaller than 1% and a fold increase of at least 20%; this was done for all four datasets (RAW264.7 native, RAW264.7 crosslinked, HeLa native and HeLa crosslinked) independently. This initial hitlist was then further refined in multiple steps: 1) PPIs were combined into two datasets, one for each cell line; 2) if a PPI passed the FC criterion in both conditions (native and crosslinked), the fdr requirement was loosened to fdr ≤ 0.05; 3) the resulting PPIs were ranked according to fdr and according to FC for each effector and each condition (native and crosslinked); 4) only PPIs that were in the top 20 for either FC or fdr were called “hit”; 5) in addition, all PPIs that passed the FC requirement, as well as the loosened fdr requirement in both conditions were called a “hit”. Output from tables from statistical analysis are in Table S2.

### Network building and GO-term analysis

Networks were built from the hits for both native and crosslinked pulldowns. Known host-host functional interactions (physical and/or functional from genomic context, high-throughput experiments, (conserved) co-expression and previous knowledge), as well as bacterial functional interactions were imported into cytoscape v3.7.2 (Shannon et al. 2003) using STRING protein query (STRING DB version 11 (Szklarczyk et al. 2019)) for the respective organism and a confidence cutoff of 0.7 (see Table S5 for functional interaction network edges of the different organisms). Using a reference list of all the proteins detected in the LC-MS/MS runs for the respective human (HeLa) or rodent (RAW264.7) host, GO-term enrichment for biological processes was performed using ClueGO version 2.5.2 with the cell line specific AP- QMS protein background as reference proteome. GO-term fusion, as well as grouping was enabled using a p-value cutoff of 0.05 after Benjamini Hochberg p-value correction. GO-terms contained in GO level 4 and 5 were searched, requiring at least 3 genes and 15% of genes per term and merging groups if at least 40% of genes and terms overlapped. The leading group term was chosen as the GO-term containing the largest number of genes.

### SDS-PAGE and Immunoblotting

For protein separation and detection, the BioRad system, and RunBlue precast gradient gels (expedeon) were used. Prior to loading on the gel, samples were diluted in Laemmli buffer (Laemmli 1970) containing 100 mM DTT and heated to 98°C for 10 min. Samples were spun down and loaded using a Hamilton syringe. SDS-PAGE was performed at a constant voltage of 150V for 50 min. For Western Blot, Immobilon-P PVDF or nitrocellulose membranes were used in a BioRad system (100V for 90 min while keeping the system cool). Subsequently, membranes were blocked in 5% milk in TBST for 1h and incubated in primary antibody diluted 1:1000 (see Table S14 for manufacturer and origin and antibody dilutions used) overnight. Membranes were washed 3 times for 5 min in TBST and subsequently incubated in secondary antibody conjugated to HRP (see Table S14) for 1h in 5% milk in TBST. After washing 3 times for 5 min, exposure using SuperSignalTM West Pico Plus chemiluminescent substrate (Thermo scientific) or Supersignal West Femto Max Sensitivity ECL onto Lucent Blue X-Ray films (advansta) or Kodak film in the dark was used to detect protein bands.

### Reciprocal pulldown validation

In order to validate PPIs identified from the AP-QMS workflow, we used a panel of 11 host target specific antibodies (see Table S14 for antibodies used). Per reaction, 50µl slurry of Protein-A beads (Thermo Fisher, cat. nr. 22811) for antibodies produced in rabbit or Protein G beads (Abcam, ab193259) for antibodies produced in mouse or rat, were washed twice in lysis buffer (0.1% Triton-X100 in PBS containing protease inhibitor). For each reaction, 3.5µl antibody were added to 100µl of washed beads in lysis buffer and incubated at room temperature for 2 hr with constant rotation in order to load the beads. The bead-antibody mixture was applied to the cleared, fresh lysate (obtained as described in the “Proteomic sample preparation section) without removing unbound antibody and incubated at 4°C for 4h. Samples were then centrifuged for 1 min at 5,000 rpm at 4°C and supernatant was decanted. The antigen-bound beads were washed 3 times in wash buffer (PBS containing 0.01% Triton-X100) by centrifuging at 5,000 rpm at 4°C for 1 min. After the final washing step, supernatants were removed and 100 µL of Laemmli buffer (Laemmli 1970) containing 100 mM DTT was added to the beads. Samples were heated to 98°C for 10 min followed by centrifugation for 1 min at 14,000 rpm. Eluates were analyzed by immunoblot (see Table S14 for antibody dilutions).

### PipB and PipB2 immunoprecipitations and mass spectrometry

HeLa adenocarcinoma epithelial cells (ATCC CCL-2) were grown in Eagle’s modified medium (Mediatech) containing 10% heat-inactivated fetal calf serum (Invitrogen) at 37°C with 5% CO_2_. Cells were seeded in 10 cm tissue-culture treated dishes and transfected with FuGENE 6® reagent (Roche) for 24 hr. Plasmid DNA was prepared using the QIAfilter Plasmid Midi kit (QIAGEN) according to the manufacturer’s instructions. For identification of PipB-specific interacting protein(s) (Fig S10), eight 10 cm tissue-culture treated dishes of HeLa cells were transfected with pEGCP-C1, pEGFP-PipB or pEGFP-PipB2. Monolayers were washed twice in cold PBS and collected by scraping into PBS. Cells were lysed on ice for 30 min in 50 mM Tris-HCl pH 7.6, 150 mM NaCl, 1mM EDTA, 0.1% Nonidet P-40 containing protease inhibitor cocktail set III and phosphatase inhibitor cocktail set II (EMD Biosciences). Samples were centrifuged at 3,000xg for 10 min at 4°C, the post-nuclear supernatant collected and precleared with Protein A agarose for 1 h at 4°C. The supernatant was collected and incubated with mouse anti-GFP clone 3E6 (Molecular Probes, Figure 6E) for 1 h at 4°C, followed by the addition of Protein A agarose. Beads were washed four times in lysis buffer and bound proteins eluted with boiling 1.5x SDS-PAGE sample buffer. Proteins were separated by SDS-PAGE on 4-15% gradient gels (BioRad) and visualized with SilverQuest Silver staining kit (Thermo). A 150 kDa band unique to the GFP-PipB immunoprecipitate was excised and sent for LC-MS/MS analysis at the Stanford University Mass Spectrometry (SUMS) Facility. For confirmation of the PipB-PDZD8 interaction under infection conditions (Fig 4B), HeLa cells seeded in 10 cm tissue-culture treated dishes were transfected with PDZD8-myc and infected with the following *S*. Typhimurium strains 12 h later: Δ*pipB* pPipB-2HA or Δ*pipB2* pPipB2-2HA at a MOI of 50 (ten 10 cm dishes per strain). At 12 h p.i., monolayers were collected and processed as described above. After 30 min lysis, samples were centrifuged at 6,000xg for 15 min at 4°C (which is sufficient to pellet intact bacteria), the supernatant collected and pre-cleared with Protein A agarose, followed by incubation with mouse anti-myc clone 4A6 agarose conjugate (EMD Millipore). Beads were washed in lysis buffer and bound proteins eluted with boiling 1.5x SDS-PAGE sample buffer. Immunoprecipitates were separated by SDS-PAGE and subject to immunoblotting with rabbit polyclonal anti-PDZD8 peptide antibodies and mouse monoclonal anti-HA.11 antibodies (Covance).

### Microscopy of F-actin and Filipin

Cells were seeded in 96-well glass bottom plates (Greiner CellContact, 30,000 cells per well for RAW264.7, 7,000 cells per well for HeLa or 3T3 fibroblasts) and infected with *S*Tm 14028s strains constitutively expressing mCherry from a plasmid (pFCcGi). After infection, cells were washed 3 times in warm PBS and fixed for 45 min at room temperature in 4% (w/v) formaldehyde (Thermo Scientific; 28908) in PBS containing 0.1% Triton-X100. Fixing solution was removed and cells were washed 3 times in cold PBS. Staining with Hoechst (2 µg/ml, Invitrogen, cat. nr. H3570) and Phalloidin ATTO-647N (30.6 µg/ml, Sigma, cat. nr. 65906) were performed in PBS for 1 h at room temperature. After staining, cells were washed 3 times in cold PBS then stored at 4°C in the dark prior to imaging.

For monitoring cholesterol trafficking, HeLa cells that were seeded one day prior at a density of 7,000 cells per well in a 96-well plate were infected as described above. 12 hr post infection, cells were washed twice in PBS and fixed in 4% (v/w) formaldehyde in PBS. After two washes in PBS, cells were stained with filipin (10 µg/ml in PBS, Sigma, cat. nr. F4767-1MG) for 30 min, and subsequently with HCS CellMask™ Deep Red Stain (Thermo, cat. nr. H32721) as described by the distributor for another 30 min. Cells were blocked for 1 hr at room temperature in PBS containing 3% Bovine Serum Albumin (Gerbu, cat. nr. 1062,0250 and 1062,9005), and subsequently incubated for 1h at RT with Alexa-488-coupled anti-LAMP1 antibody (1:500 in PBS containing 1% BSA, Abcam). Cells were washed twice in PBS and stored at 4°C in the dark.

Imaging was performed on a Nikon Eclipse Ti run with the NIS Elements software (version 4.60) at 10x or 20x magnification. Per well, 9 images (for 10x objective) or 16 images (for 20x objective) were taken at predefined, evenly spaced positions using the following filters: DAPI, FITC, Cy3, Cy5. Images were segmented using the Cell Profiler software (version 3.0.0). For segmentation, a nuclei mask was defined based on the DAPI channel. The identified objects were used to determine cell outlines from phalloidin staining (Cy5 channel). Finally, *S*Tm were identified using a fixed threshold from the Cy3 image and filtered with the cell mask. All primary and secondary objects were quantified and further analyzed according to the phenotypic readout (e.g. infection rate, bacterial load).

Quantification of co-localization was performed in ImageJ (version 1.51n). A cell mask was created by applying Ostu segmentation to the cell outline image after rolling background subtraction with a radius of 10 pixels (for phalloidin) or Huang segmentation after a rolling background subtraction with a 15 pixel radius (for cytostain). Similarly, a *Salmonella* mask was created by applying Otsu segmentation after rolling background subtraction with a 10 pixel radius to the Cy3 channel and overlaying it with a LAMP1 mask obtained through Ostu segmentation after a rolling background subtraction with a 10 pixel radius, where applicable. In order to quantify the degree of co-localization, the average intensity of phalloidin or filipin within the *Salmonella* mask was divided by the average intensity within the cell mask by applying the masks to the phalloidin or filipin images, calculating the integrated intensity and normalizing to the size of the cell or *Salmonella* mask. Random distribution and no co-localization hence yields a mean value of 1, while co-localization of *Salmonella* and phalloidin or filipin yields a value >1.

### Immunofluorescence

HeLa cells were seeded onto acid-washed coverslips in 24-well plates and transfected with EGFP-PipB for 24 h. Cells were fixed and permeabilized as described previously (Lau et al. 2019). Monolayers were incubated with primary antibodies - rabbit polyclonal anti-PDZD8 (Sigma; 1:100 dilution) and mouse anti-PDI (clone RL90, Affinity Bioreagents; 1:200 dilution) – followed by Alexa Fluor-conjugated secondary antibodies. Cells were mounted on glass slides in Mowiol. Alternatively, HeLa cells were transfected with pKozak-PDZD8-myc and infected 12 h later with invasive *S*. Typhimurium Δ*pipB* pPipB-2HA or Δ*pipB2* pPipB2-2HA bacteria at an MOI of 50 for 10 min. Invasive bacteria were prepared and infection conditions were as described previously (Klein, Powers, and Knodler 2017). Monolayers were fixed at 12 hpi, permeabilized and immunostained with the following primary antibodies - rat monoclonal anti-HA (clone 3F10, Roche: 1:250 dilution), mouse monoclonal anti-myc (clone 9B11, Cell Signaling; 1:2000 dilution), rabbit polyclonal anti-LAMP2 (kindly provided by Minoru Fukuda (Fukuda et al. 1988); 1:1000 dilution) or rabbit polyclonal anti-*Salmonella* O-antigen group B Factors 1,4,5,12 (Difco; 1:2,000 dilution) – followed by Alexa Fluor-conjugated secondary antibodies. Image acquisition was on a Zeiss LSM510 or LSM710 confocal microscope using sequential acquisition mode through an optical section of 0.25 µm in the z-axis. Images are maximum intensity projections of z-stacks.

### Yeast two-hybrid analysis

The AH109 yeast reporter strain was maintained on YPD agar plates. Transformation of AH109 cells with pGAD424- and pGBT9-based constructs by the lithium acetate method was performed following the guidelines in the Matchmaker two-hybrid system (Clontech). Double transformants were isolated on synthetic defined medium lacking leucine and tryptophan. Interaction of fusion proteins was monitored by activation of *HIS3* gene transcription following plating on medium lacking histidine, leucine and tryptophan (Mattera et al. 2003).

### Protein purification and size exclusion chromatography

Recombinant SteC and a catalytic inactive mutant of SteC (SteC-K256H) were expressed and purified as previously described (Poh et al. 2008). Purified SteC and SteC-K256H were then dialyzed in 20 mM Tris-HCl pH 7.4, 200 mM NaCl overnight at 4°C. Samples were concentrated in a 15 ml Amicon centrifuge column (Ultra 15, 3,000 NMWL cutoff - UCF900324), glycerol was then added to 10% final concentration and samples were snap-frozen and stored at -80°C. N-terminal recombinant GST fusion of human FMNL1 (1-385) was expressed from pGEX-4T1-tev-FMNL1-A1(1-385) in Rosetta(DE3) plysRare as follows. Briefly, GST fusion proteins were expressed overnight at 250 rpm at 25°C in autoinduction media (Studier 2005). Cells were harvested and lysed by sonication in lysis buffer (500 mM NaCl, 50 mM Tris/HCl (pH 7.8), 20% glycerol, 100 µg/ml lysozyme and 1x cOmplete mini EDTA-free protease inhibitors). GST-fusions were bound to pre-equilibrated Glutathione Sepharose 4B (GE; 17-0756-01) overnight at 4°C. Beads were then washed thrice with 100 mM NaCl, 50 mM Tris/HCl (pH 7.8), 10% glycerol. GST bound protein was then cleaved using biotinylated thrombin (Merck Millipore; 69672) according to the manufacturers instructions overnight at 4°C. Direct interactions between SteC and SteC-K256H with FMNL1 (1-385) were assessed by analytical gel filtration using an Akta FPLC UPC-900 equipped with a Superose 6 Increase 10/300 GL column (Merck). Typically, 500 μg of each protein was loaded onto the column equilibrated with 100 mM NaCl, 50 mM Tris/HCl (pH 7.8), 10% glycerol prior to sample injection. Complex formation was assessed by mixing equimolar amounts of each protein on ice for 5 min at 4°C prior to injection on the column. Optical density was monitored at 280 nm (UV) throughout the experiment. As a reference for molecular mass, a Bio-Rad protein standard (#1511901), covering 1.35 - 670 kDa was used. UV traces were combined and visualized in Prism v7.

### In vitro kinase assays

Purified recombinant SteC and catalytically inactive SteC-K256H (10μg each) kinases were pre-activated with kinase buffer (50 mM Tris-HCl pH 7.5, 10 mM MgCl_2_, 2 mM DTT and 50 µM ATP) for 5 min at 30°C. FMNL1 1-385 and FMNL2 2-478 were purified as previously described (Kühn et al. 2015). Next, 10 µg of the purified FMNL1 substrate was mixed with Tris-DTT buffer (50 mM Tris-HCl pH 7.5, 2 mM DTT) and added to the pre-activated kinase mix. Radiolabeled [^32^P]-γ-ATP was added to the mix and incubated for 30 min at 30°C. The reaction was stopped by the addition of 2x Laemmli buffer. Labeled proteins were resolved by SDS-PAGE, transferred to a PVDF membrane and detected by autoradiography. Proteins were visualized by Coomassie staining.

For phosphoproteomics, kinase pre-activation was achieved using 2 μg of SteC and SteC K256H kinases as described above. 8 μg of FMNL1 1-385 (FMNL1 sample) or 8 µg of FMNL2 2-478 (FMNL2 sample) or 4 µg of FMNL1 and 4 µg of FMNL2 (FMNL1+FMNL2 sample) were mixed with kinase buffer (50 mM Tris-HCl pH 7.5, 10 mM MgCl_2_, 2 mM DTT and 50 µM ATP) and added to the pre-activated kinase. All reactions were incubated at 30°C for 30 min and snap-frozen on dry ice and stored at -80°C. After thawing, HEPES pH 8.5 was added to a final concentration of 100 mM. Reduction/alkylation of cysteine residues was performed by addition of Tris(2-carboxyethyl)phosphine hydrochloride and chloroacetamide (Sigma-Aldrich) at final concentrations of 5 mM and 30 mM, respectively. Trypsin (Sigma-Aldrich) was added at a 1:25 ratio (w/w) and the samples were incubated overnight at room temperature. Samples were then desalted on stage-tips (Rappsilber, Ishihama, and Mann 2003) prepared in-house and packed with 1 mg of C18 material (ReproSil-Pur 120 C18-AQ 5 μm, Dr Maisch).

### LC-MS/MS Phosphoproteomics

Nanoflow LC-MS/MS analysis was performed by coupling an UltiMate 3000 RSLCnano LC system (Thermo Scientific) to a Fusion Orbitrap Lumos mass spectrometer (Thermo Scientific). Dried peptides were resuspended in a loading buffer consisting of 20 mM citric acid (Sigma-Aldrich) and 1% formic acid (Sigma-Aldrich). Peptides were injected, trapped and washed on a precolumn (C18 PepMap 100, 5µm, 300 µm i.d. x 5 mm, 100 Å, Thermo Scientific) for 3 min at a flow rate of 30 μL/min with 100% buffer A (0.1% formic acid in HPLC grade water). Peptides were then transferred into an analytical column (Waters nanoEase HSS C18 T3, 75 µm x 25 cm, 1.8 µm, 100 Å) before separation at a flow rate of 300 nL/min using a 45 min gradient, from 8% to 32% buffer B (0.1% formic acid, 80% acetonitrile, Sigma-Aldrich). Electrospray ionization was performed using a 2.1 kV spray voltage and a transfer capillary temperature of 275°C. The mass spectrometer was operated in data-dependent acquisition mode. Full mass spectra (m/z 300-1500) were acquired in the Orbitrap analyzer at a resolution of 60,000 with an Automated Gain Control (AGC) target value of 4e5 charges and a maximum injection time of 50 ms. The mass spectrometer was operated in Topspeed mode (maximum duty cycle time of 3 s) and precursors were sequentially selected to undergo HCD fragmentation at a normalized collision energy of 30%. The precursor intensity threshold was set to 1e5 and the dynamic exclusion to 8 seconds. MS2 spectra were acquired in the Orbitrap analyzer at a resolution of 30,000 (isolation window of 1.6 Th) with an AGC target value of 1e5 charges and a maximum injection time of 200 ms. Precursors with unassigned charge state as well as charge states of 1+ and ≥ 6+ were excluded from fragmentation.

MaxQuant software (version 1.6.2.3 (Cox and Mann 2008)) was used to process the raw data files, which were searched against a database consisting of FMNL1, FMNL2 and SteC proteins as well as commonly observed contaminants. The following parameters were used for the database search: trypsin digestion with a maximum of 3 missed cleavages, fixed carbamidomethylation of cysteine residues, variable oxidation of methionine residues as well as variable phosphorylation of serine/threonine/tyrosine residues and variable N-terminal acetylation. Mass tolerance was set to 4.5 ppm at the MS1 level and 20 ppm at the MS2 level. False discovery rate was set to 1%, the minimum peptide length to 7 residues, a score cut-off of 40 was used for modified peptides, and the match between runs option was used with a retention match time window of 2 min.

## Notes

### Competing Interest Statement

The authors have declared no competing interest.

