## Supplementary Table 4 for "Global mapping of *Salmonella enterica*-host protein-protein interactions during infection"

| **Interaction** | **Experimental settings and systems** | **References** |
| --- | --- | --- |
| PipB2-ATP1A1 | Bio-ID after ectopic expression of BirA-fusion in HeLa cells | (D'Costa et al. 2019) |
| PipB2-ATP2B1 | Bio-ID and AP-MS after ectopic expression of BirA-fusion in HeLa cells | (D'Costa et al. 2019) |
| PipB2-EPHA2 | Bio-ID after ectopic expression of BirA-fusion in HeLa cells | (D'Costa et al. 2019) |
| PipB2-GCN1 | AP-MS after ectopic expression of BirA-fusion in HeLa cells | (D'Costa et al. 2019) |
| PipB2-ITGB1 | Bio-ID after ectopic expression of BirA-fusion in HeLa cells | (D'Costa et al. 2019) |
| PipB2-KIF5B | Bio-ID after ectopic expression of BirA-fusion in HeLa cells  Co-purification of ectopically expressed effector and target in HeLa cells | (D'Costa et al. 2019;  Henry et al. 2006) |
| PipB2-KLC1 | Bio-ID after ectopic expression of BirA-fusion in HeLa cells  Co-purification of ectopically expressed effector and target in HeLa cells | (D'Costa et al. 2019; Henry et al. 2006; Schleker et al. 2012) |
| PipB2-KLC2 | Bio-ID after ectopic expression of BirA-fusion in HeLa cells  Co-purification of ectopically expressed effector and target in HeLa cells | (D'Costa et al. 2019; Henry et al. 2006; Schleker et al. 2012) |
| PipB2-LTN1 | AP-MS after ectopic expression of BirA-fusion in HeLa cells | (D'Costa et al. 2019) |
| PipB2-MARK2 | Bio-ID after ectopic expression of BirA-fusion in HeLa cells | (D'Costa et al. 2019) |
| PipB2-PDS5A | AP-MS after ectopic expression of BirA-fusion in HeLa cells | (D'Costa et al. 2019) |
| PipB2-PTPN1 | Bio-ID after ectopic expression of BirA-fusion in HeLa cells | (D'Costa et al. 2019) |
| PipB2-SLC12A2 | Bio-ID after ectopic expression of BirA-fusion in HeLa cells | (D'Costa et al. 2019) |
| PipB2-SLC1A5 | Bio-ID after ectopic expression of BirA-fusion in HeLa cells | (D'Costa et al. 2019) |
| PipB2-SLC3A2 | Bio-ID after ectopic expression of BirA-fusion in HeLa cells | (D'Costa et al. 2019) |
| PipB2-TNPO1 | AP-MS after ectopic expression of BirA-fusion in HeLa cells | (D'Costa et al. 2019) |
| SifA-GNL3 | Bio-ID after ectopic expression of BirA-fusion in HeLa cells | (D'Costa et al. 2019) |
| SifA-MATR3 | Bio-ID after ectopic expression of BirA-fusion in HeLa cells | (D'Costa et al. 2019) |
| SifA-RBM10 | AP-MS after ectopic expression of BirA-fusion in HeLa cells | (D'Costa et al. 2019) |
| SifA-ZC3HAV1 | Bio-ID after ectopic expression of BirA-fusion in HeLa cells | (D'Costa et al. 2019) |
| SseI-ACADM | AP-MS after incubation of recombinant effector in HeLa or RAW cell lysates | (Sontag et al. 2016) |
| SseI-Gm9755 | AP-MS after incubation of recombinant effector in HeLa or RAW cell lysates | (Sontag et al. 2016) |
| SseJ-RhoA | Y2H, colocalization  In vitro binding, activation upon expression in HeLa cells | (Ohlson et al. 2008; Christen et al. 2009; Auweter et al. 2011; Schleker et al. 2012) |
| SseJ-RhoB | Y2H, colocalization | (Ohlson et al. 2008) |
| SseL-OSBP | AP-MS after ectopic expression of HA-fusion in HEK-293T cells  Mechanistic follow-up in vitro using protein truncations | (Auweter et al. 2012; Auweter et al. 2011; Schleker et al. 2012) |
